# Differential Effects of Incomplete Lineage Sorting and Gene Tree Estimation Error on Gene Tree Distributions and Species Tree Inference

**DOI:** 10.64898/2026.02.21.707162

**Authors:** Nafis Tahmid, Shattik Islam Rhythm, Md Shamsuzzoha Bayzid

## Abstract

Accurate species tree inference from genome-scale data is complicated by gene tree discordance, which can arise both from biological processes such as incomplete lineage sorting (ILS) and from technical factors such as gene tree estimation error (GTEE). While both factors reduce the accuracy of summary methods, their relative impact and characteristic patterns remain poorly understood. Here, we systematically compare the effects of ILS and GTEE by simulating gene tree datasets with comparable overall discordance levels, but with discordance arising exclusively from either ILS or GTEE. Using widely employed summary methods such as ASTRAL and wQFM, we show that GTEE typically has a stronger detrimental effect on species tree accuracy than ILS, even at matched discordance levels. We further characterize the structure of gene tree distributions under these two sources of discordance and show that ILS induces a structured, constrained skew in quartet distributions, whereas GTEE generates more uniform, high-entropy noise that does not diminish with additional genes. Our case study on a widely used avian phylogenomic dataset reveals similar distributional patterns across exons, introns, and ultraconserved elements (UCEs), which differ substantially in their levels of phylogenetic signal. A quartet-based analysis of these gene trees further shows that prioritizing loci with stronger and more consistent quartet support can improve the recovery of established avian clades. Overall, these results provide an empirical framework for a nuanced understanding of how ILS and GTEE shape gene tree distributions and influence species tree inference from limited or noisy gene tree datasets.

**Significance:** Despite the growing awareness that gene tree discordance caused by incomplete lineage sorting (ILS) and gene tree estimation error (GTEE) presents a major challenge for species tree inference from multi-locus data, their relative effects remain poorly understood. Here, we systematically compare how ILS and GTEE alter gene tree distributions and affect species tree estimation. Our findings reveal that ILS and GTEE shape gene tree distributions in fundamentally different ways, which in turn differentially affect species tree inference, even when the overall amount of discordance is similar. In particular, GTEE can be substantially more detrimental than ILS and may generate patterns of noise that do not diminish simply by adding more genes. These findings show that the source, not only the amount, of gene tree discordance is important for interpreting phylogenomic data and for developing more robust species tree inference methods.

## 1 Introduction

With the availability of genome-scale data, phylogenomic analyses using multiple loci have become increasingly common. However, species tree estimation from multi-locus data sampled from throughout the whole genome is difficult because individual gene trees often disagree with one another and with the underlying species tree, a phenomenon known as gene tree discordance (Maddison 1997). A major biological cause of gene tree discordance is Incomplete Lineage Sorting (ILS) under the multi-species coalescent (MSC) (Kingman 1982; Pamilo and Nei 1998): lineages can fail to coalesce in ancestral populations (at speciation points), yielding true gene trees that differ from the species tree. By contrast, gene tree estimation error (GTEE) arises when finite sequence data, short internal branches due to rapid radiations (Xi et al. 2015; Cai et al. 2021), inaccurate alignments, and model and algorithm limitations cause the inferred gene tree to deviate from the true gene genealogy.

In the presence of gene tree discordance, standard methods for estimating species trees, such as concatenation (which combines sequence alignments from different loci into a single “supermatrix” and then computes a tree on the supermatrix) can be statistically inconsistent (Sebastien Roch and Steel 2015; Degnan et al. 2009) and produce incorrect trees with high support (L. S. Kubatko and Degnan 2007). Therefore, *summary methods* that operate by first estimating gene trees independently and then combining them to infer a species tree are becoming increasingly popular (Liu, Yu, and Edwards 2010; Mirarab, Reaz, et al. 2014; Larget et al. 2010; Mossel and Roch 2011; L. S. Kubatko, Carstens, et al. 2009; Chifman and L. Kubatko 2014; Liu, Yu, Pearl, et al. 2009; Liu and Yu 2011; Mahbub et al. 2021; Avni et al. 2015; Islam et al. 2020). Many of these summary methods are provably statistically consistent meaning that they will converge in probability to the true species tree given sufficiently large numbers of genes and sites per gene.

Despite their theoretical appeal, summary methods exhibit substantial sensitivity to gene tree estimation error (GTEE) (Bayzid and Tandy Warnow 2013; Gatesy and Springer 2013). Increasing sequence length can reduce GTEE, but long contiguous sequences are more likely to contain intralocus recombination (Hayley C. Lanier and Knowles 2012), which violates the assumption of a single underlying genealogy per locus–an essential condition for MSC-based summary approaches. This tension has contributed to a sustained debate on whether summary methods should be preferred over concatenation in practice, even when ILS is present (Gatesy and Springer 2013; Gatesy and Springer 2014; Springer and Gatesy 2016).

Due to the growing awareness, both empirical (Bayzid and Tandy Warnow 2013; Hayley C Lanier et al. 2014; Bayzid, Mirarab, et al. 2015; Blom et al. 2017; Xi et al. 2015) and theoretical (Sebastien Roch and Tandy Warnow 2015), that GTEE can substantially impair the accuracy and statistical guarantee of summary methods, designing summary methods that are robust to GTEE is key to constructing phylogenetic trees from genome-wide data. While both ILS and GTEE are recognized as major contributors to gene tree–species tree discordance and the impacts of ILS and GTEE on species tree accuracy have each been extensively studied, their relative impacts under comparable levels of discordance have not been systematically characterized. In particular, most empirical phylogenomic analyses summarize discordance as an aggregate property of the gene tree set, making it difficult to determine from such analyses alone how different sources of discordance would affect species tree inference if they acted under controlled and comparable conditions. As a result, it remains difficult to obtain a nuanced understanding of how ILS and GTEE differentially reshape gene tree topology distributions and propagate uncertainty into estimated species trees, particularly in finite-locus settings where increasing the number of genes does not necessarily eliminate gene tree estimation uncertainty. In this study, we address this gap by investigating the following two research questions (RQs).

1. RQ1: How do incomplete lineage sorting (ILS) and gene tree estimation error (GTEE) differentially affect species tree inference accuracy when they induce comparable levels of gene tree discordance?
2. address this question, we compare the effects of ILS and GTEE under controlled simulation conditions with matched “equivalent” levels of discordance. We emphasize that this study does not attempt to separate mixed biological and estimation-driven sources of discordance in empirical gene tree datasets. Instead, our objective is to understand their comparative effects by creating simulation settings in which discordance is induced exclusively by by either ILS or GTEE. This controlled design allows us to ask how ILS-induced and GTEE-induced discordance affect species tree inference when their overall discordance levels are comparable. ILS is controlled by varying effective population sizes and branch lengths under the MSC, while GTEE is regulated by altering sequence lengths. We then assess the impact of these distinct sources of discordance on species tree accuracy using several widely used summary methods.
3. RQ2: How do ILS and GTEE shape the statistical properties of gene tree topology distributions, and how do these differences influence species tree inference?
4. Here, we investigate gene tree distributions with “equivalent” discordance induced separately by ILS and GTEE across a range of discordance levels and quantify how each discordance source alters key features of the topology space and quartet support profiles in order to elucidate the mechanisms by which ILS and GTEE perturb gene tree distributions.
5. investigate these questions using extensive analyses of simulated datasets, together with analyses of a biological dataset used to examine whether patterns observed in empirical gene tree distributions are consistent with the insights obtained from controlled simulations. Our results show that ILS and GTEE perturb gene tree distributions in fundamentally different ways that can be characterized using various statistical properties of gene tree distributions. ILS produces biased and structured distributions with strong concentration around the true topology, whereas GTEE leads to more uniform, flatter, and more diffuse distributions, introducing noise that does not necessarily diminish with increasing numbers of genes. Our analyses of the avian phylogenomics dataset (Jarvis et al. 2014) reveal further important insights that are consistent with and further support these findings.

## 2 Materials and Methods

We generated datasets in a controlled simulation protocol to ensure that in any given dataset only one source of discordance was present (either ILS, which we call ILS-only, or GTEE (GTEE-only), but not both). We simulated two datasets for our study: a 15-taxon dataset containing 1000 loci and a 21-taxon dataset containing 2000 loci. Each dataset consists of 10 independent replicates, each replicate having a distinct species tree simulated using SimPhy (Mallo et al. 2016). For a particular reference species tree, we generate both GTEE-only and ILS-only conditions.

To generate datasets in which discordance arises solely from GTEE, we used the true species tree as true gene trees (indicating no ILS) for gene sequence simulation (see Figure 1), following the approach of (Cai et al. 2021). For each replicate, we used AliSim (Ly-Trong et al. 2022) to simulate alignments for the desired number of loci (1000 loci for the 15-taxon dataset and 2000 loci for the 21-taxon dataset) under three different sequence lengths: 1000, 500, and 250 base pairs. Gene trees were then inferred from each alignment using IQ-TREE 3 (Wong et al. 2026). This resulted, for each replicate, in three GTEE-only conditions: gene trees inferred from 1000 bp alignments (low-discordance), from 500 bp alignments (moderate-discordance), and from 250 bp alignments (high-discordance). Because all sequences were simulated directly from the species tree, any discordance between the inferred gene trees and the true species tree in these datasets arises exclusively from estimation error. Algorithm 1 presents the procedure for constructing the GTEE-only datasets.

**Fig. 1:**
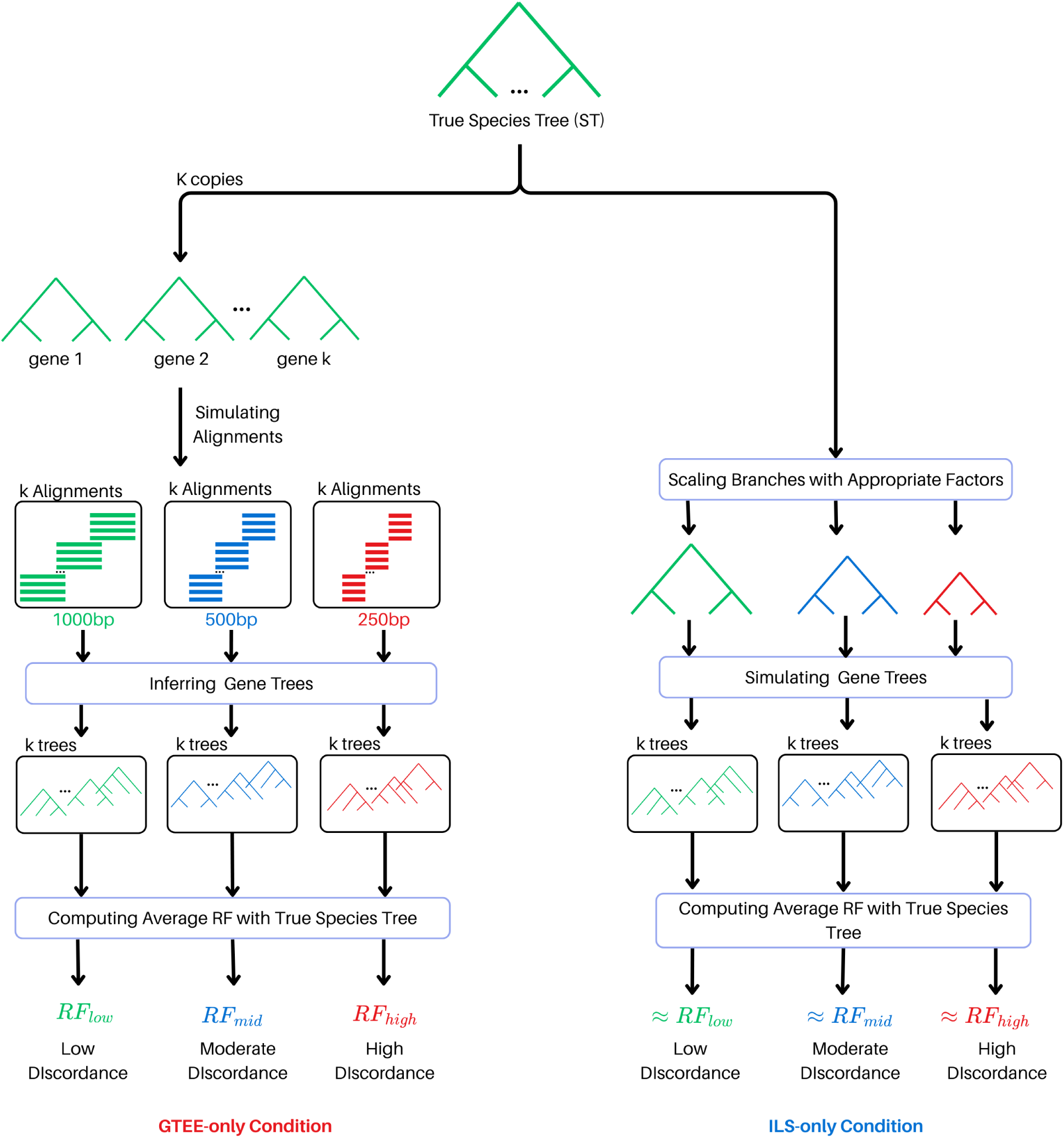
Overview of the dataset generation pipeline. Simulated true species trees (ST) were copied *k* times and treated as the true gene trees to generate sequence alignments of varying lengths (1000 bp, 500 bp, and 250 bp). Gene trees were then inferred from these simulated alignments. Variation in sequence length induced different levels of gene tree estimation error (GTEE), resulting in low, moderate, and high discordance, quantified using the average Robinson–Foulds (RF) distance between inferred gene trees and the true species tree (ST). To generate datasets with equivalent levels of discordance driven solely by incomplete lineage sorting (ILS), species tree branch lengths were scaled by appropriate factors, and gene trees were simulated directly from the scaled species trees. The scaling factors were selected to match the average RF distances observed under the corresponding GTEE-induced discordance levels (low, moderate, and high), thereby establishing three equivalent condition pairs, each comprising a GTEE-only scenario and its matched ILS-only discordance setting.

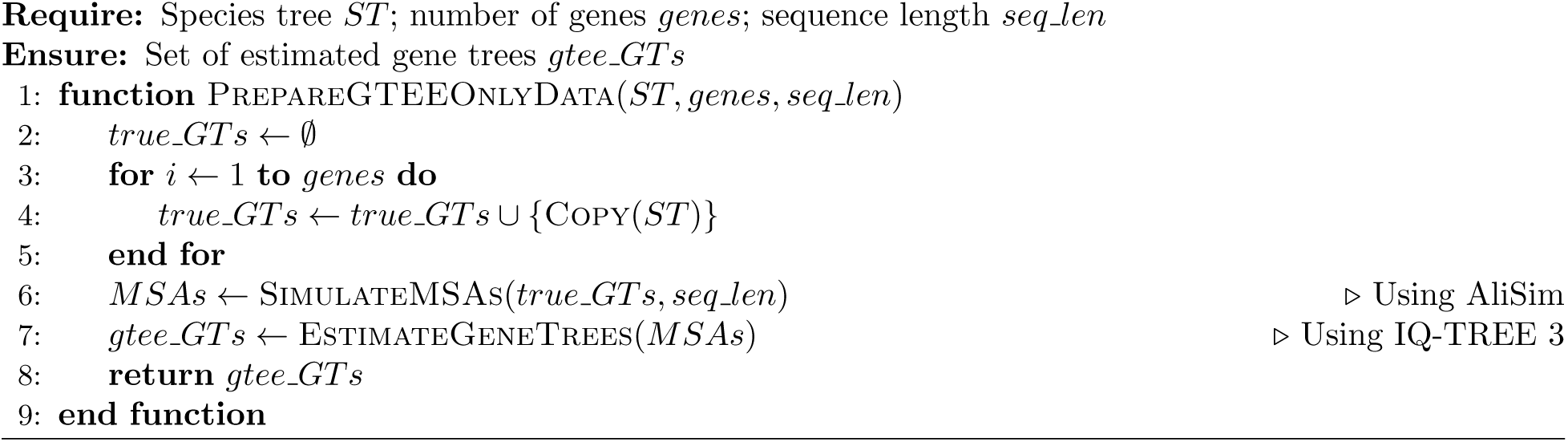

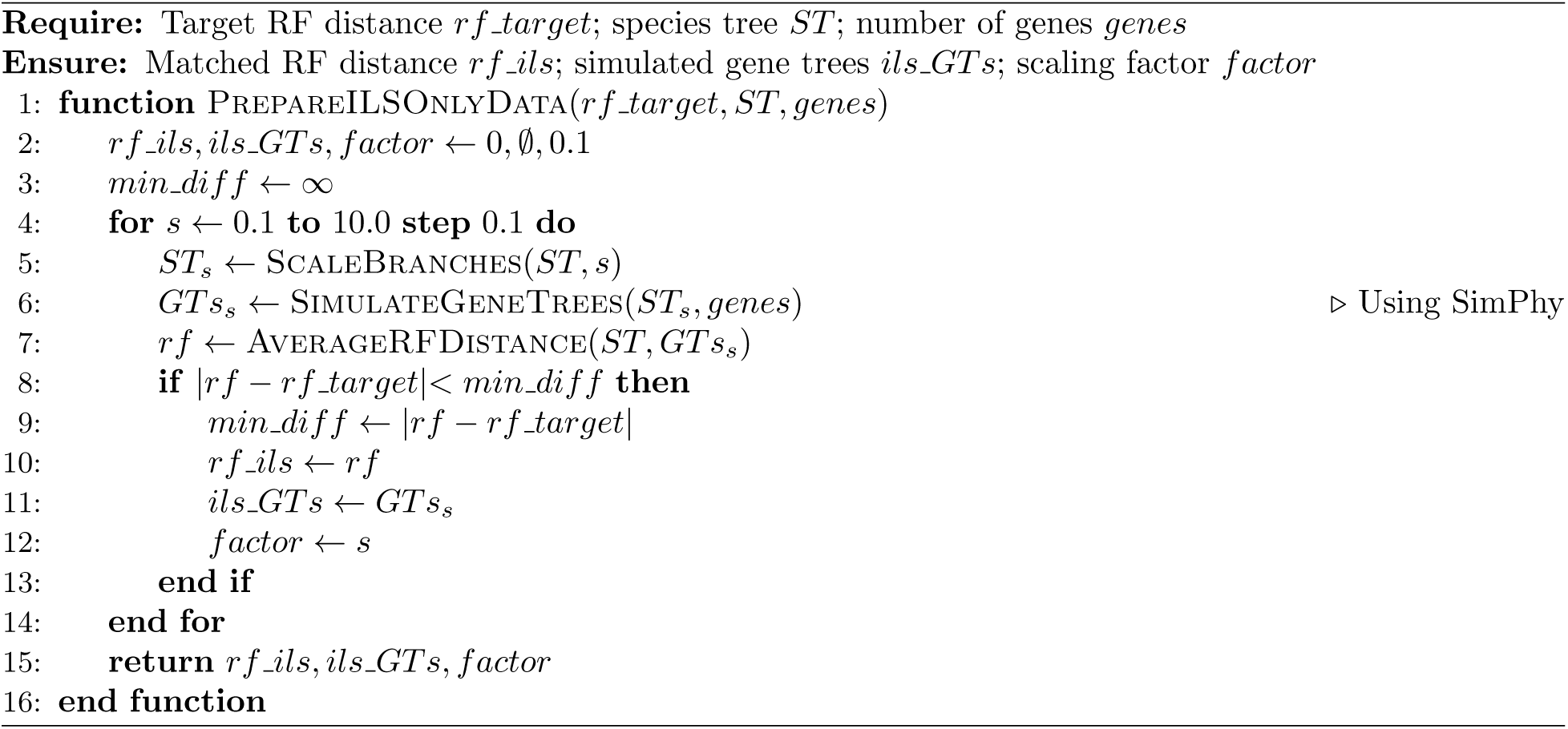

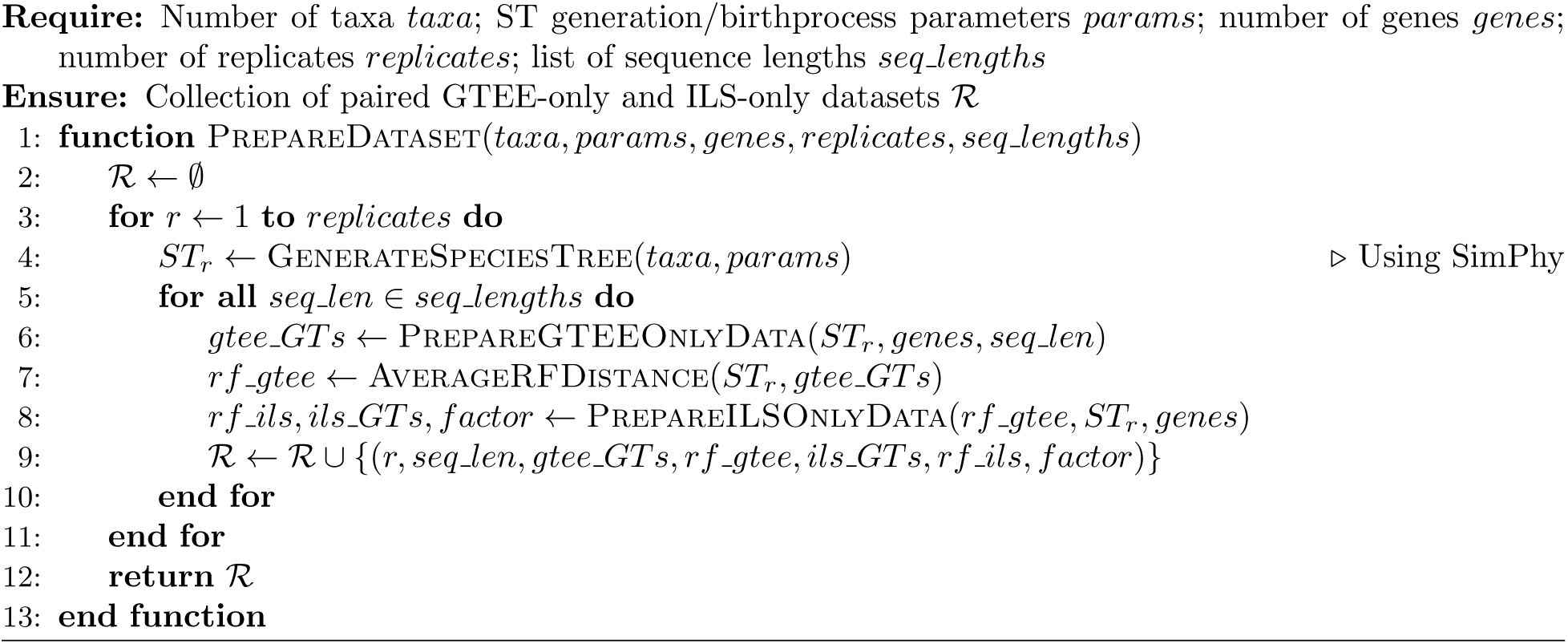

To construct ILS-only datasets with discordance levels matched to the GTEE conditions, we first quantified the level of discordance present in each GTEE dataset using the average normalized RF distance between the species tree *S* and a set *G* of gene trees. We compute the discordance levels for each replicate under each GTEE condition (low, moderate and high), and use these values as targets for constructing corresponding ILS-only datasets. For each replicate, we used SimPhy to simulate gene trees by systematically scaling the branch lengths of the same species tree used in the GTEE-only simulations. By adjusting the branch length scaling, we controlled the amount of ILS such that the resulting average RF distance between the simulated gene trees and the species tree closely matched the discordance observed under each GTEE condition (Algorithm 2). In this way, we constructed paired scenarios with comparable discordance levels in which one dataset exhibits discordance attributable solely to estimation error and the other exhibits an equivalent level of discordance attributable solely to ILS.

We invoked PrepareDataset (15*, params,* 1000, 10, [250, 500, 1000]) and PrepareDataset (21*, params,* 2000, 10, [250, 500, 1000]) to generate the 15-taxon and 21-taxon datasets, respectively, where *params* denotes the species tree simulation parameters (see Algorithms 1–3).

A schematic diagram of the data simulation workflow is shown in Figure 1. Full details of the simulation parameters, branch length scaling factors, and the resulting achieved discordance levels and quartet scores for the datasets are provided in Supplementary Section 1.

### 2.1 Species tree estimation methods

We investigated the impact of ILS and GTEE on different types of methods. Under the Maximizing Quartet Consistency (MQC) criterion (Snir and Rao 2012; Reaz et al. 2014; Mirarab, Reaz, et al. 2014), we considered ASTRAL (Mirarab, Reaz, et al. 2014), a widely used and highly accurate statistically consistent summary method, and wQFM (Mahbub et al. 2021), a leading quartet-amalgamation algorithm that also optimizes MQC and has been shown to match or exceed ASTRAL’s accuracy. We also considered the minimize deep coalescence (MDC) criterion (Maddison 1997; C. V. Than and Nakhleh 2009; Yang and Warnow 2011), which seeks the species tree that minimizes the number of deep coalescence events required for a given collection of gene trees. Phylonet (C. V. Than, Ruths, et al. 2008) is a popular tool for estimating species trees under MDC. Although it has been shown to be statistically inconsistent under the multispecies coalescent model (C. V Than and Rosenberg 2011), this is not agnostic to the gene tree heterogeneity as it takes into account the specific nature of the way incomplete lineage sorting occurs. Finally, we included Greedy Consensus (Kozlov et al. 2019), a topology-only approach that is agnostic to the source of discordance.

### 2.2 Measurements

We compared the estimated trees (on simulated datasets) with the model species tree using normalized Robinson-Foulds (RF) distance to measure the tree error. For the biological data set, we compared the estimated species trees with the existing literature and biological beliefs. We also performed different experiments based on the quartet scores (the number of quartets in the gene trees that agree with a species tree) of the trees estimated by different methods.

## 3 Results

In this section, we present the results related to the research questions (RQs) explored in this study.

### 3.1 RQ1: Effects of the source of discordance on species tree estimation under controlled simulations

To examine the relative impact of ILS and GTEE on species tree inference, we established equivalence between ILS-only and GTEE-only model conditions by matching the overall levels of gene tree discordance, as described in Section 2. We then inferred species trees using different species tree estimation methods on datasets with varying levels of discordance and different numbers of genes. Figure 2(a) shows the accuracies of different species tree estimation methods under equivalent ILS-only and GTEE-only model conditions for the 15-taxon dataset. Panels (b), (c), and (d) in Figure 2 and Supplementary Table 3 confirm that our experimental setup achieves comparable levels of gene tree discordance between the ILS-only and GTEE-only scenarios across varying numbers of gene trees.

**Fig. 2:**
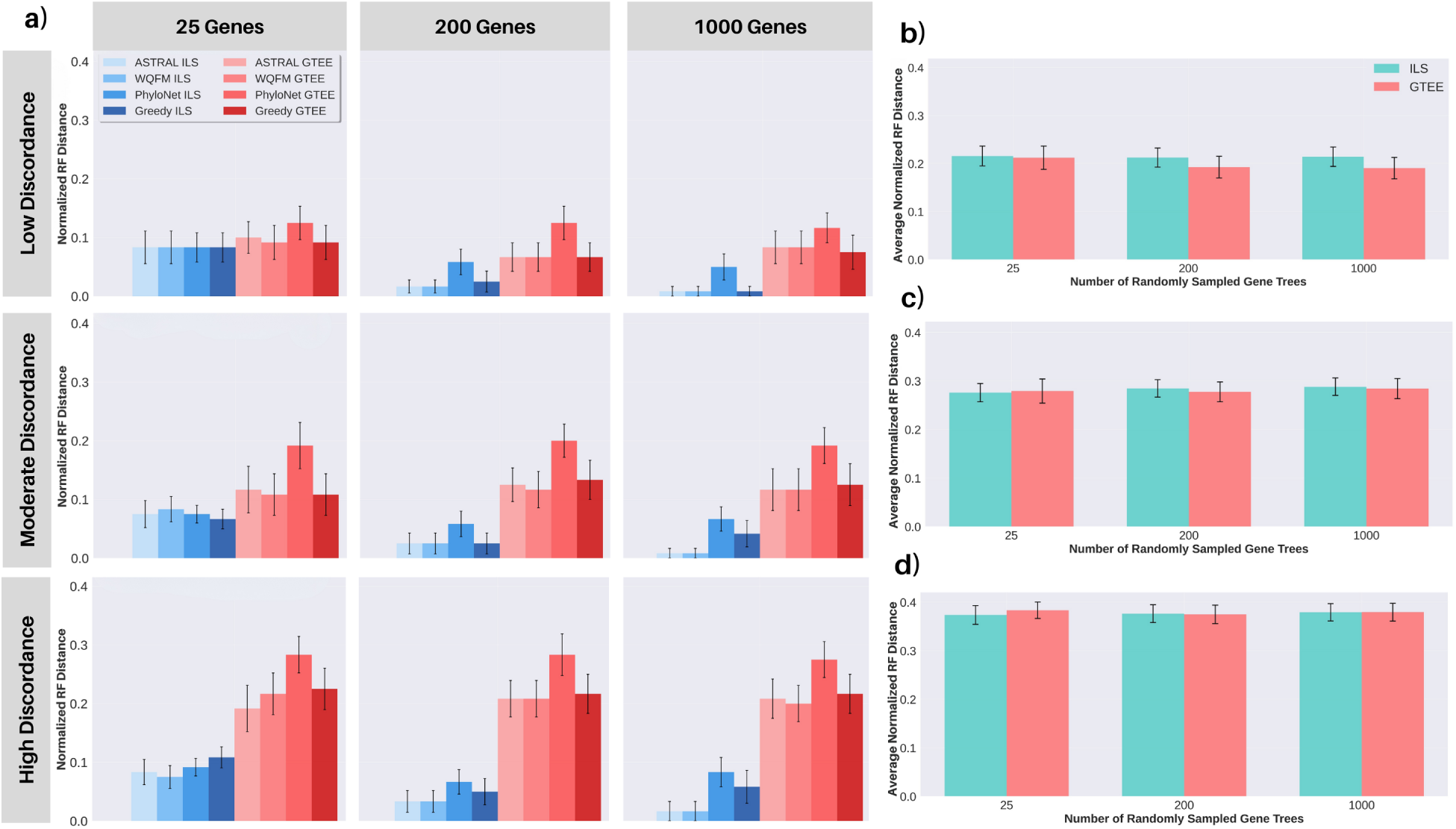
(a) Average normalized Robinson–Foulds (RF) distance between estimated and true species trees for the 15-taxon dataset under matched ILS-only and GTEE-only discordance conditions across low, moderate, and high discordance levels and gene set sizes of 25, 200, and 1000. Species trees were inferred using ASTRAL, wQFM, PhyloNet, and Greedy consensus. Bars show averages across replicates and whiskers indicate standard error. (b–d) Average normalized RF distance between input gene trees and the true species tree under (b) low, (c) moderate, and (d) high discordance levels across gene set sizes (25, 200, 1000) for the matched ILS-only and GTEE-only conditions.

The general expectation that species tree accuracy decreases with increasing levels of gene tree discordance holds across all model conditions. However, for the same level of discordance, species tree inference is consistently less accurate under the GTEE-only condition (red bars in Figure 2(a)) than under the ILS-only condition (blue bars in Figure 2(a)). This trend becomes more pronounced as the level of discordance increases. For instance, with 200 genes and low discordance, the average normalized RF rates of different methods range between 1.7-5.8% for the ILS-only datasets, compared to 6.7-12.5% for the GTEE-only datasets, representing an average increase of about *∼*5% (Supplementary Table 4). At higher discordance, such as in the high-discordance model with 200 genes, the difference between the ILS-only and GTEE-only conditions becomes much larger, with RF rates of around 3.3-6.7% and 20.8-28.3% under the ILS-only and GTEE-only conditions, respectively. These results clearly demonstrate the differential impact of the two sources of gene tree discordance on species tree accuracy and indicate that GTEE is substantially more detrimental than ILS on the methods tested.

We next examine the impact of ILS-only and GTEE-only conditions as the number of gene trees increases. As expected, for ILS-only datasets at a given level of discordance, the RF rate decreases as the number of gene trees increases. This effect is particularly notable for methods such as ASTRAL and wQFM, which are statistically consistent and thus can benefit from increased gene sampling. In contrast, Phylonet and greedy consensus, which lack statistical consistency, do not show notable improvements with increasing numbers of genes. On the other hand, in the GTEE-only setting, a very different pattern was observed. Unlike the ILS-only case, species tree accuracy does not improve noticeably as more gene trees are sampled. For instance, on the high-discordance datasets, RF rates remain almost unchanged even when the number of genes is increased from 25 to 1000, even for statistically consistent methods such as ASTRAL and wQFM.

This suggests that the noise introduced by GTEE has a particularly harmful effect on species tree inference, likely because it disrupts the assumptions underlying the statistical consistency of summary methods. Unlike discordance generated by ILS, GTEE-driven discordance may produce error patterns that do not average out simply by increasing the number of genes. Therefore, when discordance is driven primarily by GTEE, adding more gene trees alone is insufficient to guarantee improved species tree accuracy unless sequence lengths are also increased, thereby reducing GTEE and the resulting discordance. Comparable inference trends are observed in the 21-taxon dataset under equivalent conditions (Supplementary Figure 1, Table 5).

Overall, these results highlight that although both ILS and GTEE introduce gene tree discordance, their impacts on species tree inference are fundamentally different. These findings not only support existing knowledge that summary methods are highly sensitive to GTEE, but also demonstrate that discordance arising from ILS versus GTEE affects species tree estimation methods in distinct ways, even when the overall level of gene tree discordance is matched.

### 3.2 RQ2: Investigate how ILS and GTEE perturb gene tree distributions

We performed a series of experiments to understand how ILS and GTEE perturb gene tree distributions, even when both induce comparable overall levels of discordance from the true species tree. The goal is to investigate systematic differences in the patterns of discordance that may explain downstream differences in species tree inference accuracy.

First, we examined the prevalence of true species tree quartets within the gene tree distributions under ILS- and GTEE-only model conditions with equivalent discordance levels. Because quartets are statistically consistent estimators of the species tree under ILS, it is important to understand how ILS and GTEE perturb the quartet distributions induced by gene trees. For each set of four taxa *{a, b, c, d}*, we let *q^∗^* denote the quartet topology present in the true species tree and computed its frequency in the gene tree distributions under both the ILS- and the GTEE-only conditions. The difference Δ = freq_ILS_(*q^∗^*) *−* freq_GTEE_(*q^∗^*) was calculated for all such 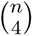 true species tree quartets, where *n* denote the number of taxa. Figure 3 (a–c) shows the sorted Δ values across the three discordance levels (low, moderate and high) for a representative replicate (R4 in Supplementary Table 3, 6) of the 15-taxon dataset (all replicates: Supplementary Figure 7). Across all settings, a clear and consistent pattern emerges: the vast majority of true quartets are more frequent under the ILS-only condition than under the GTEE-only condition, as reflected by Δ *>* 0 for most quartets as well as the mean Δ being above zero in most cases. These confirm that ILS-only gene tree sets contain true quartets at comparatively higher frequencies than their GTEE-only counterparts. In addition, the true quartets are more frequently the dominant topology under ILS-only conditions than under GTEE-only conditions across different levels of discordance (Supplementary Table 6). Here, dominant topology refers to the most frequently observed quartet topology in the gene trees among the three possible quartet resolutions of any four-taxa set. Under GTEE, the proportion of true quartets that remain the dominant topology generally decreases with increasing levels of discordance. Therefore, as alignment length decreases and GTEE (discordance) increases, the true quartet signal generally becomes weaker, whereas under the matched ILS conditions, true quartets more frequently remain the dominant topology, indicating greater preservation of the underlying biological signal across different levels of discordance.

**Fig. 3:**
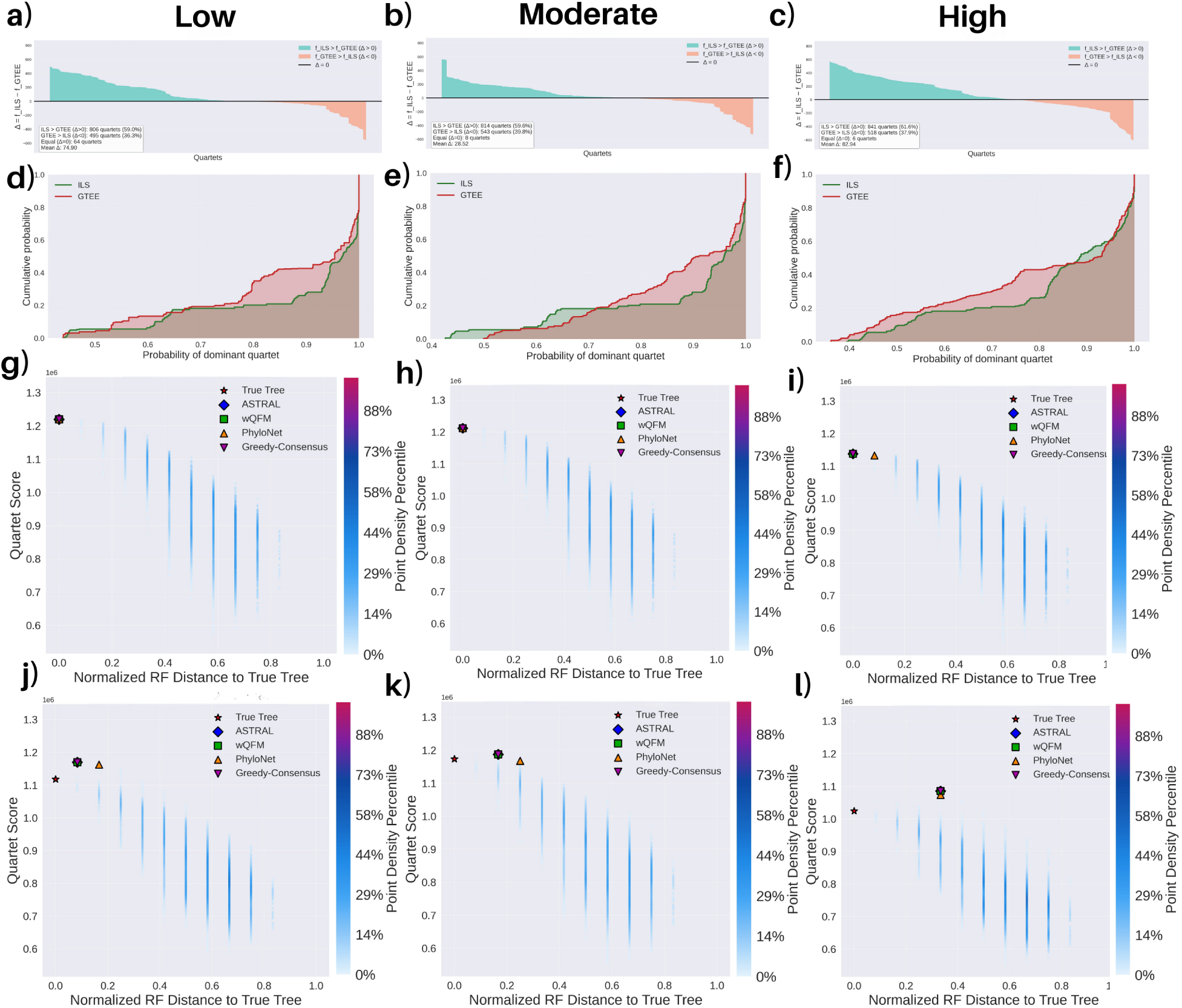
Quartet distribution and species-tree neighborhood characteristics under matched ILS-only and GTEE-only discordance conditions for a representative replicate of the 15-taxon dataset under under low, moderate, and high discordance levels. (a–c) Sorted differences in true species-tree quartet (*q^∗^*) frequencies (Δ = freq_ILS_(*q^∗^*) *−* freq_GTEE_(*q^∗^*)) across all quartets 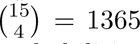. (d–f) Empirical cumulative distribution functions of dominant quartet probabilities for matched ILS-only and GTEE-only conditions. (g–i) Quartet score landscapes of 20,000 SPR-sampled neighboring species trees plotted against normalized RF distance to the true species tree under ILS-only conditions. Here, we also show ASTRAL, wQFM, PhyloNet, and GC-estimated trees. (j–l) Quartet score landscapes under GTEE-only conditions. We color the data points with a color gradient which varies continuously from light blue to red with increasing density of data points (species trees).

Second, we examined how ILS and GTEE influence quartet-level support across gene tree distributions. Given a true species tree *S^∗^* and for a set of four taxa *{a, b, c, d}*, let *q^∗^* be the quartet topology induced by *S^∗^* with an internal branch of length *τ >* 0 (coalescent units), and *q*_1_*, q*_2_ the other two alternative topologies. Under the multi-species coalescent model, the probability distribution is:

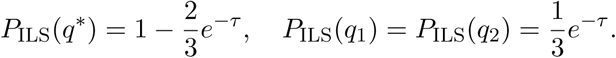

Therefore, for any *τ >* 0, the distribution is strictly skewed with the true topology having the highest probability (*P*_ILS_(*q^∗^*) *> P*_ILS_(*q*_1_) = *P*_ILS_(*q*_2_)) and its Shannon entropy *H*_ILS_(*τ*) is strictly less than log 3 (the entropy of a uniform distribution on three outcomes). Thus, ILS is expected to induce a structured and biased quartet distribution. In contrast, GTEE is expected to make the three alternate quartet topologies more similar in frequency. Thus, under GTEE, the quartet probability distribution is expected to be more diffuse. To investigate whether this theoretical understanding is reflected in the simulated datasets, we show (Figure 3, panels d–f for replicate R4; all replicates: Supplementary Figures 3–4) the empirical cumulative distribution function (ECDF) of dominant quartet probabilities across all 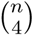 quartets, where *n* is the number of taxa. Here, if a curve for one dataset lies below the curve for another dataset, then the former strictly improves on the latter with respect to the support of the dominant quartets. For our representative replicate, under ILS, the ECDFs remain relatively flat across the low-to-mid probability range and then rise sharply at high probabilities (approximately above 0.85), indicating that a large fraction of dominant quartets have high probabilities. In the low and moderate discordance settings, about 75% of dominant quartets exceed a probability of 0.9, and even under high discordance about 75% exceed 0.8. In contrast, under GTEE-only conditions, the ECDFs increase more gradually, indicating that a large fraction of the most frequent topology fails to achieve strong dominance and the probability gap between the dominant and alternative resolutions is reduced. Under low and moderate discordance, only about 55% and 50% of dominant quartets, respectively, exceed a probability of 0.9, and under high discordance only about 55% exceed 0.8.

We further summarized these patterns by computing the skewness of the dominant quartet probability distribution. Besides, for each four-taxon set, we computed the entropy, 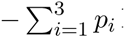 and dispersion, 1*−*max(*p*_1_*, p*_2_*, p*_3_), where *p*_1_*, p*_2_*, p*_3_ are the probabilities of the three possible quartet resolutions of a four-taxon set. Entropy and dispersion were then averaged across all 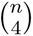 four-taxon sets. Across discordance levels, the summary statistics generally indicate more concentrated and asymmetric quartet support under ILS and flatter, more dispersed support under GTEE (Supplementary Table 6).

Finally, we examined how these distributional differences affect the species tree space. We sampled 20,000 distinct species-tree topologies in the local neighborhood of the true species tree using subtree-prune-and-regraft (SPR) rearrangements. To obtain a set of nearest neighbors, we enumerated candidate topologies by breadth-first search (BFS) over the SPR graph: we first generated all unique topologies reachable by exactly one SPR move from the true tree, and only if additional topologies were needed did we proceed to topologies reachable by two SPR moves, then three, and so on. During this search, branch lengths were ignored (topologies only), and duplicate topologies were removed. Each sampled topology was then evaluated by its normalized RF distance to the true species tree and by its quartet score (the number of gene-tree quartets consistent with that species tree). We then plotted the quartet scores of these trees against their corresponding RF rates (Figure 3(g)–(l)). We identified pseudo-terraces (Farah et al. 2021), regions of tree space containing multiple distinct tree topologies (i.e., different RF rates) with identical quartet scores, under both ILS- and GTEE-only conditions. However, the placement of the true species tree within this landscape differs between the two conditions. Under ILS-only conditions (Figure 3(g)–(i)), the true species tree generally attained the highest quartet score relative to the sampled neighboring trees. In contrast, under GTEE-only conditions (Figure 3(j)–(l)), substantially more neighboring trees achieved higher quartet scores than the true species tree, with this tendency generally becoming more pronounced as discordance increases. This overall pattern is observed across replicates (Supplementary Table 6), suggesting that, in the presence of GTEE, methods that maximize quartet score (e.g., ASTRAL and wQFM) may be more likely to favor a topology with a higher quartet score than the true species tree, potentially leading them away from the true species tree. This observation is aligned with prior studies (Farah et al. 2021). Similar trends are observed for the 21-taxon dataset, with replicate R2 shown as a representative example in Supplementary Figure 1 and results for all replicates provided in Supplementary Figures 5, 6, and 8 and Supplementary Table 7. Therefore, GTEE does not merely introduce stochastic noise into gene tree estimation, but can distort the quartet score surface in ways that may mislead quartet-based species tree inference methods. By contrast, even when ILS induces comparable levels of discordance, the true species tree generally has a higher quartet score than other candidate species trees. This supports the statistical consistency of quartet-based methods given large numbers of error-free gene trees.

### 3.3 Case study: avian biological dataset

Our earlier analyses used simulated datasets to establish equivalent discordance conditions under ILS and GTEE and examined their effects on gene tree distributions and species tree inference. Although controlled matching of discordance levels is not possible for real biological datasets, evaluating whether trends similar to those observed in our simulated datasets are also present in empirical data provides an important test for the practical relevance of our findings. To this end, we analyzed the avian phylogenomics dataset (Jarvis et al. 2014) comprising 48 taxa and 144,446 genes, including exons, introns and ultraconserved elements (UCEs). Birds are known to have undergone rapid radiations, leading to short internal branches and high levels of ILS. In addition, many loci in this dataset are relatively short and contain limited phylogenetic signal. As a result, this dataset likely contains substantial contributions from both ILS and GTEE, making it well suited for our study.

The dataset contains 8,295 exon, 2,516 intron, and 3,679 UCE loci, each with different length distributions (Figure 4(a)). Exon loci are the shortest on average (mean: 1,634.1 bp, range: 99–15,819 bp) and show a strongly right-skewed distribution, with most loci being short and only a few much longer. This suggests that many exon loci contain limited phylogenetic signal and are therefore more prone to GTEE. UCE loci are of intermediate length (mean: 2,508.7 bp, range: 180–3,561 bp) and have a relatively narrow distribution. In contrast, intron loci are much longer (mean: 7,762.4 bp, range: 82–77,090 bp) and show the widest spread, including many very long alignments.

**Fig. 4:**
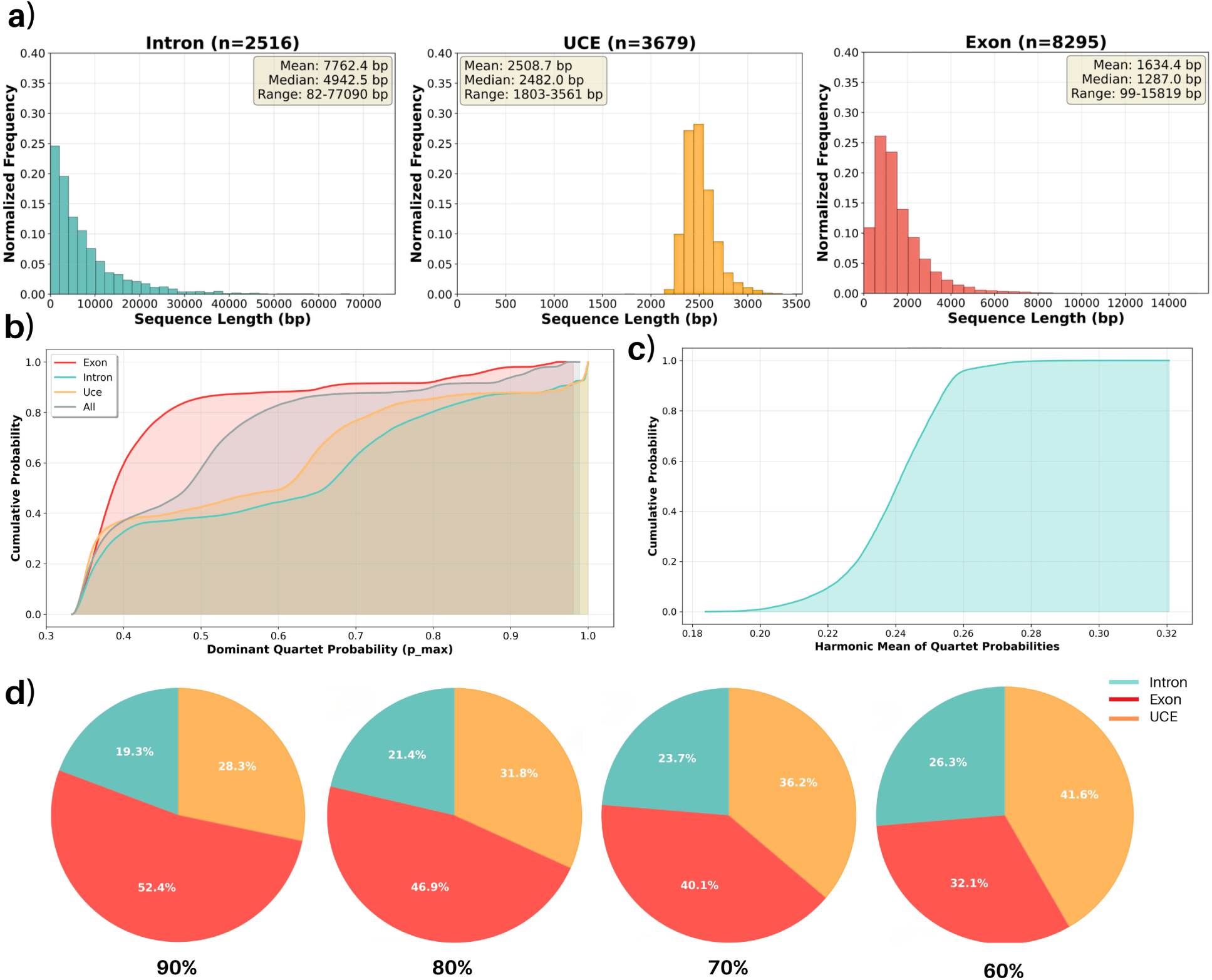
Sequence length and quartet-based characteristics of gene trees in the avian phylogenomic dataset. (a) Sequence length distributions for introns (n = 2516), UCEs (n = 3679), and exons (n = 8295), shown as normalized-frequency histograms with insets reporting mean, median, and range (bp). (b) Empirical cumulative distributions (ECDFs) of dominant quartet probabilities computed for intron, UCE, exon, and combined (all) gene sets. (c) ECDF of scores computed for all gene trees, where each gene tree is assigned a score defined as the harmonic mean of quartet probabilities estimated from the quartet distribution induced by the combined (all) set of gene trees. (d) Composition of intron, UCE, and exon loci among the top 90%, 80%, 70%, and 60% of genes trees (ranked by harmonic-mean quartet score).

We next computed dominant quartet probability distributions separately for exon, intron, and UCE gene trees (as in Section 3.2) and plotted their ECDFs (Figure 4(b)). The three locus types show clearly different patterns, reflecting differences in phylogenetic signal. The exon ECDF rises steeply, with about 60% of quartets below probability 0.40, nearly 85% below 0.50, and around 90% below 0.70, which indicate weak and diffuse support for a large portion of the dominant quartets. In contrast, the intron ECDF increases more gradually: only about 35% of quartets fall below 0.40 and roughly 60% are below 0.70, suggesting that a larger fraction of quartets attain higher probabilities and therefore provide stronger phylogenetic signal. The UCE curve shows intermediate behavior, with about 40% of quartets below 0.40 and around 75% below 0.70. These differences align with the sequence length distributions of the three locus types, since longer sequences generally provide more signal and lower gene tree estimation error. When we compute the ECDF using all 14,446 gene trees together, the curve closely follows the exon pattern across most of the range. This is expected because exons dominate the dataset (57.1%), compared to 17.4% introns and 25.5% UCEs, and therefore largely determine the overall distribution.

Overall, these patterns match those observed in the simulated datasets (Section 3.2): loci expected to have higher GTEE show flatter ECDF curves due to weaker quartet support, while loci with lower estimation error show steeper curves. This agreement between biological and simulated results supports the conclusion that sequence length, through its effect on GTEE, plays a key role in shaping quartet structure even in biological datasets with substantial ILS.

To further characterize the gene tree distributions, we quantified skewness and entropy of the dominant quartet probability distributions for each locus type. Exon loci exhibit high skewness (2.28) and the largest entropy (0.34). The presence of strong positive skewness indicates a distribution in which probability mass is concentrated near lower-support quartets, with only a small fraction achieving high dominance. The high entropy further suggests substantial uncertainty in quartet resolution, consistent with the expectation that shorter alignments amplify GTEE and lead to more similar support among the three possible quartet topologies, thereby weakening the dominance of the most frequent topology. UCEs display moderate level of skewness (0.56) and entropy (0.28). These values point to a flatter and more balanced probability distribution compared to exons. Introns exhibit the lowest skewness (0.21) and entropy (0.26), a combination characteristic of which suggests that dominant quartet probabilities are more evenly distributed toward higher support values, reflecting stronger phylogenetic signal.

Next, to quantify the phylogenetic reliability of gene trees, we ranked all 144,446 gene trees using a quartet-based scoring framework. For each set of four taxa, the probability *P* (*q_i_*) of a specific quartet topology *q_i_* is defined as the frequency of that topology in the set of gene trees divided by the total frequency of the three alternative topologies. Given a gene tree *g_i_* containing *n_i_* taxa, we evaluate all 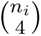 quartets present in that tree and define its score as the harmonic mean of their probabilities: 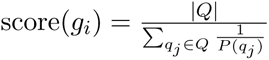, where *Q* denotes the set of quartets induced by gene tree *g_i_* and 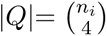. The harmonic mean is particularly suitable in this context because it strongly penalizes low-support quartets. A gene tree cannot achieve a high score simply by containing a few strongly supported quartets; instead, it must maintain consistent support across most quartets. This property ensures that higher scores indicate stronger phylogenetic signal and more reliable gene tree estimates. Next, we investigated the ECDF of the harmonic mean scores of all gene trees (Figure 4(c)). It shows a sharp rise around a score of approximately 0.23, with roughly 95% of gene trees having scores less than or equal to 0.26. This pattern indicates that the majority of loci carry limited phylogenetic signal and only a small fraction of genes achieve higher scores. Such a distribution is expected in clades shaped by rapid radiation, where short divergence amplify both ILS and estimation error (Liu, Yu, Wu, et al. 2023).

We next examined how the composition of gene categories changes when we restrict the analysis to higher-scoring loci. In the full dataset, exons account for 57.1% of gene trees, introns 17.4%, and UCEs 25.5%. But as we retain only the top 90%, 80%, 70%, or 60% of gene trees based on their scores, the proportion of exon loci decreases rapidly (Figure 4(d)). In contrast, introns and UCEs become relatively more prevalent. This pattern indicates that exon gene trees tend to receive lower reliability scores, consistent with higher GTEE due to their shorter sequence lengths. In contrast, intron loci increase steadily in representation, reflecting their stronger phylogenetic signal. Under the most stringent cutoff (top 60%), about 90.6% of introns are retained, and they make up 26.3% of the selected gene trees. These retention patterns closely follow the earlier distributional results: loci with stronger phylogenetic signal (lower skewness and lower entropy) are preferentially retained, whereas loci with greater uncertainty in quartet resolution are progressively filtered out.

Next, we examined the species trees estimated on different subsets of avian gene trees. Using the filtered gene tree sets defined by harmonic-mean quartet scores, we inferred species trees using ASTRAL and wQFM. Trees inferred from the top 90% (least stringent) and top 60% (most stringent) gene sets are shown in Figure 5 and were compared against the avian reference phylogeny estimated using MP-EST on binned gene trees (Jarvis et al. 2014; Mirarab, Bayzid, et al. 2014). We also included trees from prior studies (Mahbub et al. 2021) inferred on exons, introns, and UCEs using ASTRAL and wQFM. The recovery of established avian clades by ASTRAL and wQFM using different subsets of gene trees is summarized in Table 1. This analysis is aligned with our previous analyses, with exon-only trees recovering the fewest clades, UCE-only trees showing improved recovery (exceeding exon and all-gene analyses), and intron-only trees achieving the highest clade recovery. For ASTRAL, trees inferred from the top 60% gene set exhibited improved clade recovery relative to both the all-gene and top 90% analyses. Notably, ASTRAL-60 uniquely recovered *Otidimorphae* and *Cursorimorphae*, clades absent in the ASTRAL-all and ASTRAL-90 trees. A similar pattern was observed for wQFM: trees inferred from the top 90% and 60% gene sets displayed comparable performance and improved upon the all-gene analysis, including recovery of *Cursorimorphae*, which was absent from the wQFM-all tree. Overall, our findings on the biological gene tree distributions exhibit trends similar to those observed in our simulations.

**Fig. 5:**
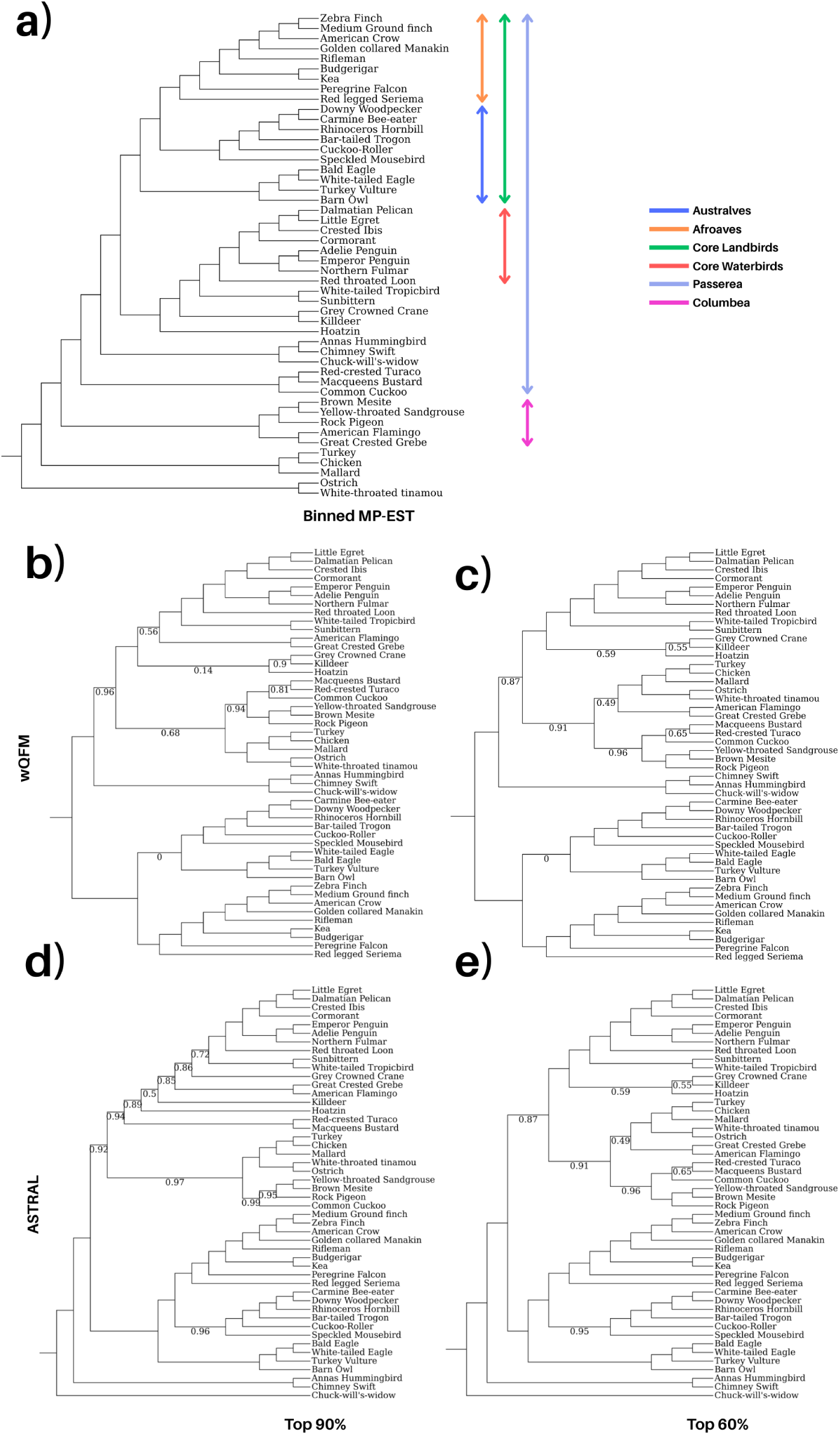
Species trees inferred from ranked gene subsets in the avian dataset. (a) Reference phylogeny (Jarvis et al. 2014) showing major avian clades. (b–c) wQFM trees from the top 90% and top 60% gene subsets, respectively. (d–e) ASTRAL trees from the top 90% and top 60% gene subsets, respectively. All branch support values are 100% except where noted. Species trees from the combined (all) gene set and from intron, UCE, and exon gene sets using ASTRAL and wQFM were estimated in prior studies (Mahbub et al. 2021).

**Table 1:**
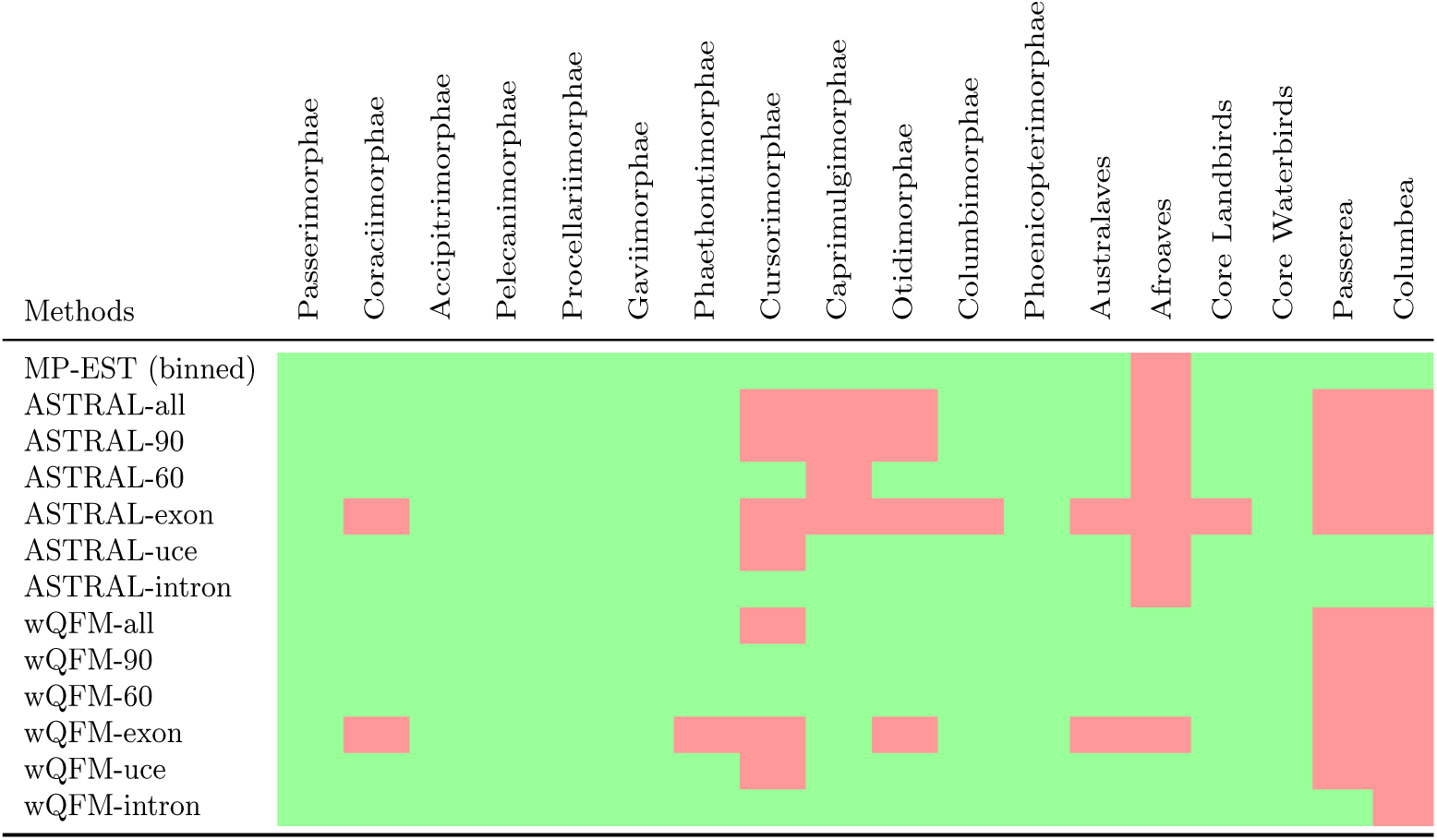
Recovery of clades in analyses of different subsets of the avian datasets. Grids show the presence (green) or absence (red) of clades. We show the results for ASTRAL and wQFM on different gene sets: all genes, the top 60% or 90% ranked by quartet score (Section 3.3), or locus-specific subsets (introngs, exons and UCEs). We also show the reference tree from (Jarvis et al. 2014) estimated by MP-EST on binned gene (Mirarab, Bayzid, et al. 2014) trees

## 4 Conclusion

In this study, we systematically compared the effects of ILS- and GTEE-induced discordance on gene tree topology distributions and species tree inference under controlled conditions with comparable overall levels of discordance. It is well known that both ILS and GTEE complicate species tree estimation, and that methods developed under ILS-based models can be highly sensitive to gene tree estimation error (Bayzid and Tandy Warnow 2013). In particular, GTEE-induced discordance may be interpreted as additional ILS because these methods generally operate on estimated gene trees without explicitly distinguishing the source of discordance, and estimated species trees may attain higher quartet scores than the true species tree, potentially misleading quartet-based methods such as ASTRAL and wQFM (Farah et al. 2021). However, the differential effects of ILS and GTEE under controlled and matched levels of discordance have remained relatively poorly understood. Moreover, little is known about how these two sources of discordance differently perturb the underlying gene tree topology distribution.

Our results clearly show that ILS and GTEE shape gene tree distributions in fundamentally different ways, which in turn differentially affect species tree inference. Under ILS-only conditions, quartet support remains structured and biased toward true quartet topologies, and increasing the number of genes improves accuracy as expected under statistical consistency. In contrast, GTEE produces flatter and more diffuse quartet distributions, and thus weakens the dominance of true quartets. As a result, even statistically consistent methods may fail to improve with additional genes when discordance is driven primarily by estimation error.

Analyses of the avian phylogenomic dataset further reveal trends similar to those observed in our simulations. Loci with stronger phylogenetic signal tend to receive higher scores under our quartet-based scoring framework and may help improve species tree estimation. These empirical patterns are consistent with the distributional differences observed in our simulations and demonstrate the potential utility of characterizing gene tree distributions for species tree inference.

Taken together, our findings indicate that ILS and GTEE have differential impacts and possibly identifiable patterns in the resulting gene tree distributions. This opens interesting research avenues to design statistical tests for indentifying the reasons for gene tree discrodance. Moreover, understanding these differences between ILS- and GTEE-induced discordance can guide the development of more robust species tree methods and motivate strategies for identifying or correcting error-prone loci (Hasan et al. 2025). Such insights are particularly important for genome-scale datasets, where the number of loci is large but gene tree discordance and uncertainty remain substantial.

## Data Availability

The datasets used to perform this study are available at https://doi.org/10.5281/zenodo. 18705938.

## Additional files

**Supplementary material SM1:** Supplementary results and additional methodological information.

## References

1. Avni, Eliran, Reuven Cohen, and Sagi Snir (2015). “Weighted quartets phylogenetics”. In: Systematic Biology 64.2, pp. 233–242.

2. Bayzid, Md Shamsuzzoha, Siavash Mirarab, Bastien Boussau, and Tandy Warnow (2015). “Weighted statistical binning: enabling statistically consistent genome-scale phylogenetic analyses”. In: PLOS One 10.6.

3. Bayzid, Md Shamsuzzoha and Tandy Warnow (2013). “Naive binning improves phylogenomic analyses”. In: Bioinformatics 29.18, pp. 2277–2284.

4. Blom, Mozes PK, Jason G Bragg, Sally Potter, and Craig Moritz (2017). “Accounting for uncertainty in gene tree estimation: summary-coalescent species tree inference in a challenging radiation of Australian lizards”. In: Systematic Biology 66.3, pp. 352–366.

5. Cai, Liming et al. (2021). “The perfect storm: gene tree estimation error, incomplete lineage sorting, and ancient gene flow explain the most recalcitrant ancient angiosperm clade, Malpighiales”. In: Systematic Biology 70.3, pp. 491–507.

6. Chifman, Julia and Laura Kubatko (2014). “Quartet from SNP data under the coalescent model”. In: Bioinformatics 30.23, pp. 3317–3324.

7. Degnan, J H, M DeGiorgio, D Bryant, and N A Rosenberg (2009). “Properties of consensus methods for inferring species trees from gene trees”. In: Systematic Biology 58, pp. 35–54.

8. Farah, Ishrat Tanzila, Muktadirul Islam, Kazi Tasnim Zinat, Atif Hasan Rahman, and Shamsuzzoha Bayzid (2021). “Species tree estimation from gene trees by minimizing deep coalescence and maximizing quartet consistency: a comparative study and the presence of pseudo species tree terraces”. In: Systematic Biology 70.6, pp. 1213–1231.

9. Gatesy, John and Mark S Springer (2013). “Concatenation versus coalescence versus “concatalescence””. In: Proceedings of the National Academy of Sciences 110.13, E1179–E1179.

10. Gatesy, John and Mark S Springer (2014). “Phylogenetic analysis at deep timescales: unreliable gene trees, bypassed hidden support, and the coalescence/concatalescence conundrum”. In: Molecular Phylogenetics and Evolution 80, pp. 231–266.

11. Hasan, Navid Bin, Sohaib, and Md Shamsuzzoha Bayzid (2025). “QT-WEAVER: Correcting quartet distribution improves phylogenomic analyses despite gene tree estimation error”. In: RECOMB International Workshop on Comparative Genomics. Springer, pp. 150–156.

12. Islam, Mazharul, Kowshika Sarker, Trisha Das, Rezwana Reaz, and Md Shamsuzzoha Bayzid (2020). “STELAR: A statistically consistent coalescent-based species tree estimation method by maximizing triplet consistency”. In: BMC Genomics 21.1, pp. 1–13.

13. Jarvis, Erich D et al. (2014). “Whole-genome analyses resolve early branches in the tree of life of modern birds”. In: Science 346.6215, pp. 1320–1331.

14. Kingman, J F C (1982). “The coalescent”. In: Stochastic Processes and their Applications 13, pp. 235–248.

15. Kozlov, Alexey M, Diego Darriba, Tomas Flouri, Benoit Morel, and Alexandros Stamatakis (2019). “RAxML-NG: a fast, scalable and user-friendly tool for maximum likelihood phylogenetic inference”. In: Bioinformatics 35.21, pp. 4453–4455.

16. Kubatko, L S, B C Carstens, and L L Knowles (2009). “Stem: Species tree estimation using maximum likelihood for gene trees under coalescence”. In: Bioinformatics 25, pp. 971–973.

17. Kubatko, L S and J H Degnan (2007). “Inconsistency of phylogenetic estimates from concatenated data under coalescence”. In: Systematic Biology 56, p. 17.

18. Lanier, Hayley C, Huateng Huang, and L Lacey Knowles (2014). “How low can you go? The effects of mutation rate on the accuracy of species-tree estimation”. In: Molecular Phylogenetics and Evolution 70, pp. 112–119.

19. Lanier, Hayley C. and L. Lacey Knowles (Jan. 2012). “Is Recombination a Problem for Species-Tree Analyses?” In: Systematic Biology 61.4, pp. 691–701. issn: 1063-5157. doi: 10.1093/sysbio/ syr128. eprint: https://academic.oup.com/sysbio/article-pdf/61/4/691/24563814/ syr128.pdf. url: 10.1093/sysbio/syr128.

20. Larget, B, S K Kotha, C N Dewey, and C Ane (2010). “BUCKy: Gene tree/species tree reconciliation with the Bayesian concordance analysis”. In: Bioinformatics 26.22, pp. 2910–2911.

21. Liu, Liang and Lili Yu (2011). “Estimating species trees from unrooted gene trees”. In: Systematic Biology 60.5, pp. 661–667. doi: 10.1093/sysbio/syr027.

22. Liu, Liang, Lili Yu, and Scott V Edwards (2010). “A maximum pseudo-likelihood approach for estimating species trees under the coalescent model”. In: BMC Evolutionary Biology 10:302.

23. Liu, Liang, Lili Yu, Dennis K Pearl, and Scott V Edwards (2009). “Estimating species phylogenies using coalescence times among sequences”. In: Systematic Biology 58.5, pp. 468–477.

24. Liu, Liang, Lili Yu, Shaoyuan Wu, et al. (2023). “Short branch attraction in phylogenomic inference under the multispecies coalescent”. In: Frontiers in Ecology and Evolution 11, p. 1134764.

25. Maddison, W. P. (1997). “Gene trees in species trees”. In: Systematic Biology 46, pp. 523–536.

26. Mahbub, Mahim, Zahin Wahab, Rezwana Reaz, M. Saifur Rahman, and Md Shamsuzzoha Bayzid (Nov. 2021). “wQFM: highly accurate genome-scale species tree estimation from weighted quartets”. In: Bioinformatics 37.21, pp. 3734–3743. issn: 1367-4803. doi: 10.1093/bioinformatics/btab428.

27. Mallo, Diego, Leonardo de Oliveira Martins, and David Posada (2016). “SimPhy: phylogenomic simulation of gene, locus, and species trees”. In: Systematic Biology 65.2, pp. 334–344.

28. Mirarab, Siavash, Md Shamsuzzoha Bayzid, Bastien Boussau, and Tandy Warnow (2014). “Statistical binning enables an accurate coalescent-based estimation of the avian tree”. In: Science 346.6215, p. 1250463.

29. Mirarab, Siavash, Rezwana Reaz, et al. (2014). “ASTRAL: genome-scale coalescent-based species tree estimation”. In: Bioinformatics 30.17, pp. i541–i548.

30. Mossel, E and S Roch (2011). “Incomplete lineage sorting: consistent phylogeny estimation from multiple loci”. In: IEEE/ACM Transactions on Computational Biology and Bioinformatics 7.1, pp. 166–171.

31. Pamilo, P. and M. Nei (1998). “Relationship between gene trees and species trees”. In: Molecular Biology and Evolution 5, pp. 568–583.

32. Reaz, Rezwana, Md Shamsuzzoha Bayzid, and M Sohel Rahman (2014). “Accurate phylogenetic tree reconstruction from quartets: A heuristic approach”. In: PLOS One 9.8, e104008.

33. Roch, Sebastien and Mike Steel (2015). “Likelihood-based tree reconstruction on a concatenation of aligned sequence data sets can be statistically inconsistent”. In: Theoretical Population Biology 100, pp. 56–62.

34. Roch, Sebastien and Tandy Warnow (2015). “On the robustness to gene tree estimation error (or lack thereof) of coalescent-based species tree methods”. In: Systematic Biology 64.4, pp. 663–676.

35. Snir, Sagi and Satish Rao (2012). “Quartet MaxCut: a fast algorithm for amalgamating quartet trees”. In: Molecular Phylogenetics and Evolution 62.1, pp. 1–8.

36. Springer, Mark S and John Gatesy (2016). “The gene tree delusion”. In: Molecular Phylogenetics and Evolution 94, pp. 1–33.

37. Than, C. V and N A Rosenberg (2011). “Consistency properties of species tree inference by minimizing deep coalescences”. In: Journal of Computational Biology 18, pp. 1–15.

38. Than, C. V. and L. Nakhleh (2009). “Species Tree Inference by Minimizing Deep Coalescences”. In: PLOS Computational Biology 5.9.

39. Than, C. V., D. Ruths, and L. Nakhleh (2008). “PhyloNet: A Software Package for Analyzing and Reconstructing Reticulate Evolutionary Relationships”. In: BMC Bioinformatics 9, p. 322.

40. Ly-Trong, Nhan, Suha Naser-Khdour, Robert Lanfear, and Bui Quang Minh (2022). “AliSim: a fast and versatile phylogenetic sequence simulator for the genomic era”. In: Molecular Biology and Evolution 39.5, msac092.

41. Wong, Thomas KF et al. (2026). “IQ-TREE 3: phylogenomic inference software using complex evolutionary models”. In: Molecular Biology and Evolution 43.5, msag117.

42. Xi, Zhenxiang, Liang Liu, and Charles C Davis (2015). “Genes with minimal phylogenetic information are problematic for coalescent analyses when gene tree estimation is biased”. In: Molecular Phylogenetics and Evolution 92, pp. 63–71.

43. Yang, J and T Warnow (2011). “Fast and accurate methods for phylogenomic analyses”. In: BMC Bioinformatics. to appear.

