## Supplementary Material for "Differential Effects of Incomplete Lineage Sorting and Gene Tree Estimation Error on Gene Tree Distributions and Species Tree Inference"

#### 1 Dataset generation: overall pipeline and parameters used

Table 1: SimPhy [1] simulation parameters and their values used for generating the 15 and 21 taxa simulated datasets

| Parameter | Flag | Value | Description |
|---|---|---|---|
| Number of replicates | -RS | 10 | Number of independent simulation replicates to perform |
| Species tree leaves | -SL | 15/21 | Number of taxa (species) in the species tree (fixed value) |
| Species tree birth rate | -SB | 0.0001 | Birth rate for species tree generation (fixed at 0.0001) |
| Species tree death rate | -SD | 0.0 | Death rate for species tree generation (fixed at 0.0, pure birth process) |
| Species tree sample | -SP | 8000 | Population size parameter for species tree (fixed at 8000) |
| Locus tree replicates | -RL | 1000/2000 | Number of gene trees to simulate per species tree (fixed value) |
| Locus tree birth rate | -LB | 0.0 | Duplication rate (fixed at 0.0) |
| Locus tree death rate | -LD | 0.0 | Loss rate (fixed at 0.0) |
| Locus tree transfer rate | -LT | 0.0 | Horizontal gene transfer rate (fixed at 0.0, no transfer) |

Listing 1: Command used to simulate trees using SimPhy [1].

```
./simphy -RS <replicates> -SL <taxa> -SB <birth_rate> -SD <death_rate> -SP <population> -RL <genes> -
    LB <duplication_rate> -LD <loss_rate> -LT <gene_transfer_rate> -CS <seed> -O <output_folder>
```

Listing 2: Command used to simulate trees from a specific species tree using SimPhy [1].

```
./simphy -S <species_tree> -SB <birth_rate> -SD <death_rate> -SP <population> -RL <genes> -LB <
    duplication_rate> -LD <loss_rate> -LT <gene_transfer_rate> -CS <seed> -O <output_folder>
```

Listing 3: Command used to simulate MSAs using AliSim [2].

```
iqtree3 --alisim <output_prefix> -t <tree_path> -m GTR{<GTR_rate_parameters>}+F{<
    base_frequency_parameters>} --seqtype DNA --length <seq_length> --seed <seed>
```

Table 2: AliSim [2] simulation parameters and their values used for generating simulated gene sequence alignments. Branches of true species trees simulated using SimPhy [1] were scaled by a factor .0001 (empirically determined) prior to AliSim simulation to keep the branch lengths in suitable range for AliSim.

| **Parameter** | **Flag** | **Value** | **Description** |
|---|---|---|---|
| Substitution model | -m | GTR{`gtr`}+F{`freq`} | General Time Reversible model with:<br>• `gtr`: 6 rate parameters (TC, TA, TG, CA, CG, AG) sampled from Dirichlet(16, 3, 5, 5, 6, 15)<br>• `freq`: 4 base frequencies (A, C, G, T) sampled from Dirichlet(36, 26, 28, 32)<br>The parameters were taken from [4] |
| Sequence type | –seqtype | DNA | Specifies DNA sequence simulation |
| Sequence length | –length | 250/500/1000 | Length of simulated sequences (number of sites) |

Listing 4: Command used to estimate gene trees from simulated MSAs using IQ-TREE 3 [3]. Note that, the GTR model parameters used for simulation are not exposed while estimating gene trees.

```
iqtree3 -s <msa_file> --prefix <output_prefix> -m GTR+F --seed <seed> -quiet
```

Table 3: Calibrated branch length (BL) scaling factors, average RF distances, and average quartet scores (proportion of consistent quartets) for ILS and GTEE scenarios with matched discordance levels for the simulated 21- and 15-taxa datasets.

| #Taxa | Replicate | 1000bp BL scaling | 1000bp ILS (RF) | 1000bp GTEE (RF) | 1000bp ILS (QS) | 1000bp GTEE (QS) | 500bp BL scaling | 500bp ILS (RF) | 500bp GTEE (RF) | 500bp ILS (QS) | 500bp GTEE (QS) | 250bp BL scaling | 250bp ILS (RF) | 250bp GTEE (RF) | 250bp ILS (QS) | 250bp GTEE (QS) |
|---|---|---|---|---|---|---|---|---|---|---|---|---|---|---|---|---|
| 21 | R1 | 10.0 | 0.197 | 0.192 | 0.956 | 0.932 | 7.2 | 0.249 | 0.251 | 0.901 | 0.866 | 5.8 | 0.296 | 0.295 | 0.840 | 0.779 |
| | R2 | 2.8 | 0.270 | 0.268 | 0.870 | 0.809 | 1.8 | 0.386 | 0.387 | 0.821 | 0.758 | 1.4 | 0.454 | 0.457 | 0.786 | 0.732 |
| | R3 | 9.8 | 0.165 | 0.165 | 0.940 | 0.924 | 4.5 | 0.305 | 0.305 | 0.867 | 0.830 | 3.8 | 0.350 | 0.347 | 0.841 | 0.825 |
| | R4 | 4.1 | 0.158 | 0.157 | 0.910 | 0.881 | 2.2 | 0.277 | 0.280 | 0.886 | 0.863 | 1.4 | 0.402 | 0.407 | 0.863 | 0.842 |
| | R5 | 1.2 | 0.413 | 0.406 | 0.855 | 0.751 | 0.8 | 0.530 | 0.518 | 0.805 | 0.742 | 0.6 | 0.610 | 0.616 | 0.765 | 0.742 |
| | R6 | 10.0 | 0.124 | 0.048 | 0.958 | 0.972 | 10.0 | 0.124 | 0.118 | 0.958 | 0.902 | 5.1 | 0.216 | 0.219 | 0.915 | 0.893 |
| | R7 | 3.7 | 0.208 | 0.207 | 0.925 | 0.896 | 3.5 | 0.213 | 0.213 | 0.924 | 0.880 | 1.6 | 0.386 | 0.382 | 0.856 | 0.767 |
| | R8 | 2.4 | 0.268 | 0.268 | 0.900 | 0.851 | 2.5 | 0.258 | 0.261 | 0.904 | 0.872 | 1.4 | 0.417 | 0.409 | 0.832 | 0.785 |
| | R9 | 5.2 | 0.185 | 0.186 | 0.918 | 0.840 | 2.9 | 0.282 | 0.280 | 0.877 | 0.790 | 1.1 | 0.501 | 0.492 | 0.768 | 0.692 |
| | R10 | 3.7 | 0.243 | 0.241 | 0.923 | 0.911 | 2.7 | 0.319 | 0.321 | 0.887 | 0.830 | 1.6 | 0.485 | 0.481 | 0.803 | 0.763 |
| 15 | R1 | 10.0 | 0.219 | 0.126 | 0.888 | 0.843 | 10.0 | 0.219 | 0.177 | 0.840 | 0.777 | 4.9 | 0.406 | 0.404 | 0.743 | 0.667 |
| | R2 | 5.5 | 0.121 | 0.120 | 0.959 | 0.919 | 4.1 | 0.177 | 0.173 | 0.933 | 0.893 | 2.3 | 0.309 | 0.313 | 0.876 | 0.843 |
| | R3 | 10.0 | 0.172 | 0.161 | 0.933 | 0.902 | 4.1 | 0.299 | 0.301 | 0.890 | 0.804 | 2.1 | 0.407 | 0.410 | 0.838 | 0.785 |
| | R4 | 4.0 | 0.286 | 0.287 | 0.894 | 0.819 | 3.8 | 0.298 | 0.301 | 0.888 | 0.859 | 2.4 | 0.406 | 0.407 | 0.833 | 0.750 |
| | R5 | 4.2 | 0.272 | 0.275 | 0.934 | 0.955 | 2.8 | 0.348 | 0.349 | 0.934 | 0.934 | 1.5 | 0.503 | 0.509 | 0.861 | 0.773 |
| | R6 | 8.8 | 0.191 | 0.190 | 0.938 | 0.896 | 3.1 | 0.336 | 0.337 | 0.863 | 0.812 | 2.6 | 0.373 | 0.372 | 0.839 | 0.789 |
| | R7 | 10.0 | 0.116 | 0.106 | 0.941 | 0.939 | 4.9 | 0.251 | 0.249 | 0.863 | 0.881 | 3.7 | 0.320 | 0.318 | 0.827 | 0.820 |
| | R8 | 4.8 | 0.284 | 0.282 | 0.828 | 0.827 | 3.6 | 0.342 | 0.346 | 0.790 | 0.816 | 3.3 | 0.362 | 0.363 | 0.781 | 0.777 |
| | R9 | 4.6 | 0.216 | 0.216 | 0.895 | 0.886 | 2.7 | 0.329 | 0.328 | 0.838 | 0.794 | 2.3 | 0.376 | 0.369 | 0.813 | 0.807 |
| | R10 | 10.0 | 0.264 | 0.140 | 0.929 | 0.945 | 8.7 | 0.278 | 0.278 | 0.924 | 0.889 | 5.9 | 0.328 | 0.328 | 0.907 | 0.844 |

#### 2 Supplementary Plots and Figures

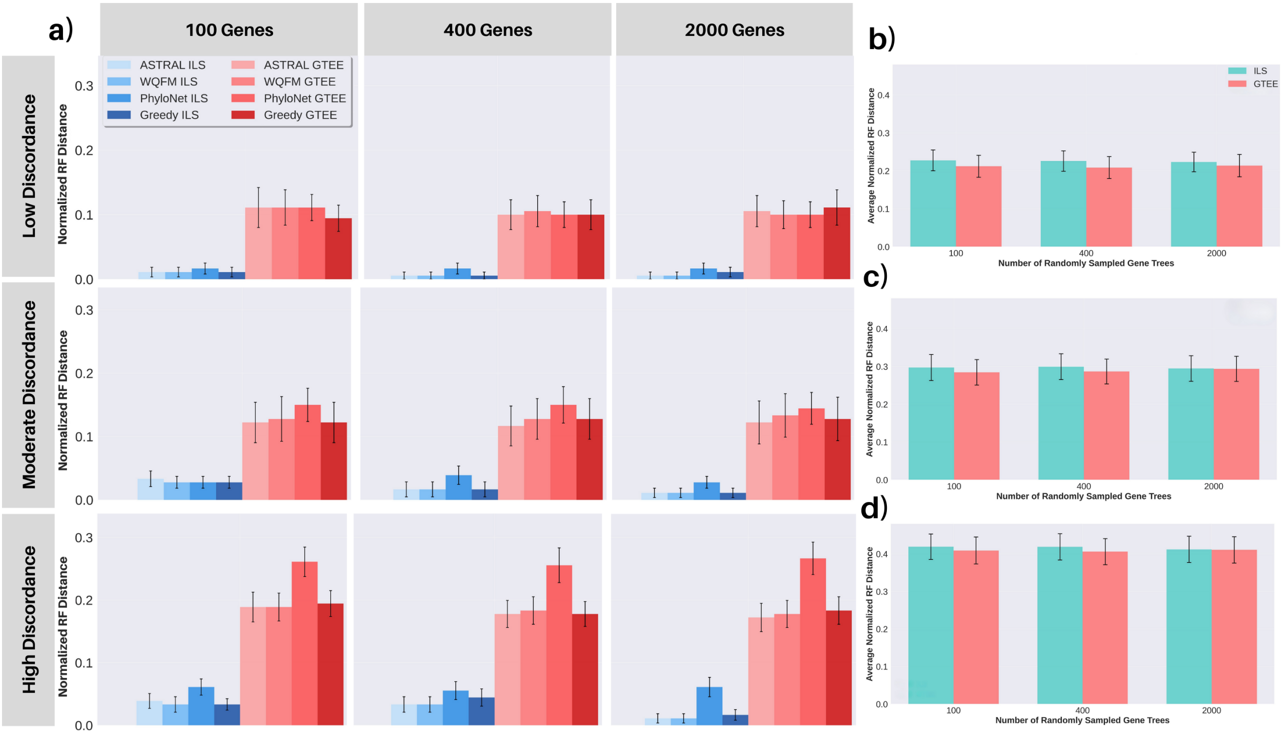


Figure 1: (a) Average normalized Robinson–Foulds (RF) distance between estimated and true species trees for the 21-taxon dataset under matched ILS-only and GTEE-only discordance conditions across low, moderate, and high discordance levels and gene set sizes of 100, 400, and 2000. Species trees were inferred using ASTRAL, wQFM, PhyloNet, and Greedy consensus. Bars show averages across replicates and whiskers indicate standard error. (b–d) Average normalized RF distance between input gene trees and the true species tree under (b) low, (c) moderate, and (d) high matched discordance across gene set sizes (25, 200, 1000) for the corresponding ILS-only and GTEE-only conditions.

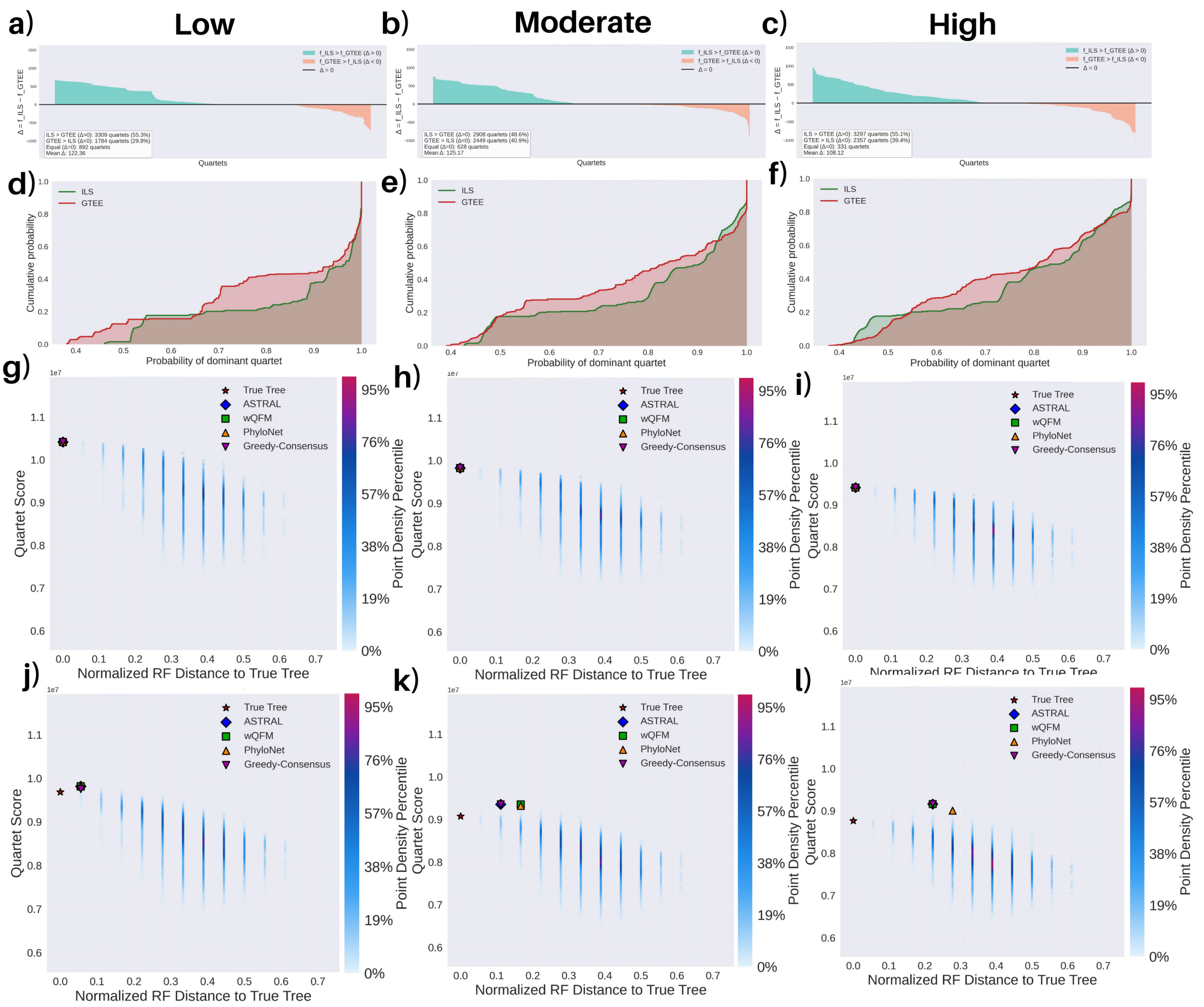


Figure 2: Quartet distribution and species-tree neighborhood characteristics under matched ILS-only and GTEE-only discordance conditions for a representative replicate (R2) of the 21-taxon dataset. (a–c) Sorted differences in true species-tree quartet ($q^*$) frequencies ($\Delta = \text{freq}_{\text{ILS}}(q^*) - \text{freq}_{\text{GTEE}}(q^*)$) across all quartets (total $\binom{21}{4} = 5985$ quartets) under low, moderate, and high discordance (for all replicates: see Supplementary Figure 10). (d–f) Empirical cumulative distribution functions of dominant quartet probabilities for the corresponding discordance levels under matched ILS-only and GTEE-only conditions (for all replicates: see Supplementary Figure 7-8). (g–i) Quartet score landscapes of 20,000 SPR-sampled neighboring species trees plotted against normalized RF distance to the true species tree under ILS-only conditions for low, moderate, and high discordance; markers denote the true species tree and species trees inferred using ASTRAL, wQFM, PhyloNet, and Greedy consensus. (j–l) Corresponding quartet score landscapes under GTEE-only conditions.

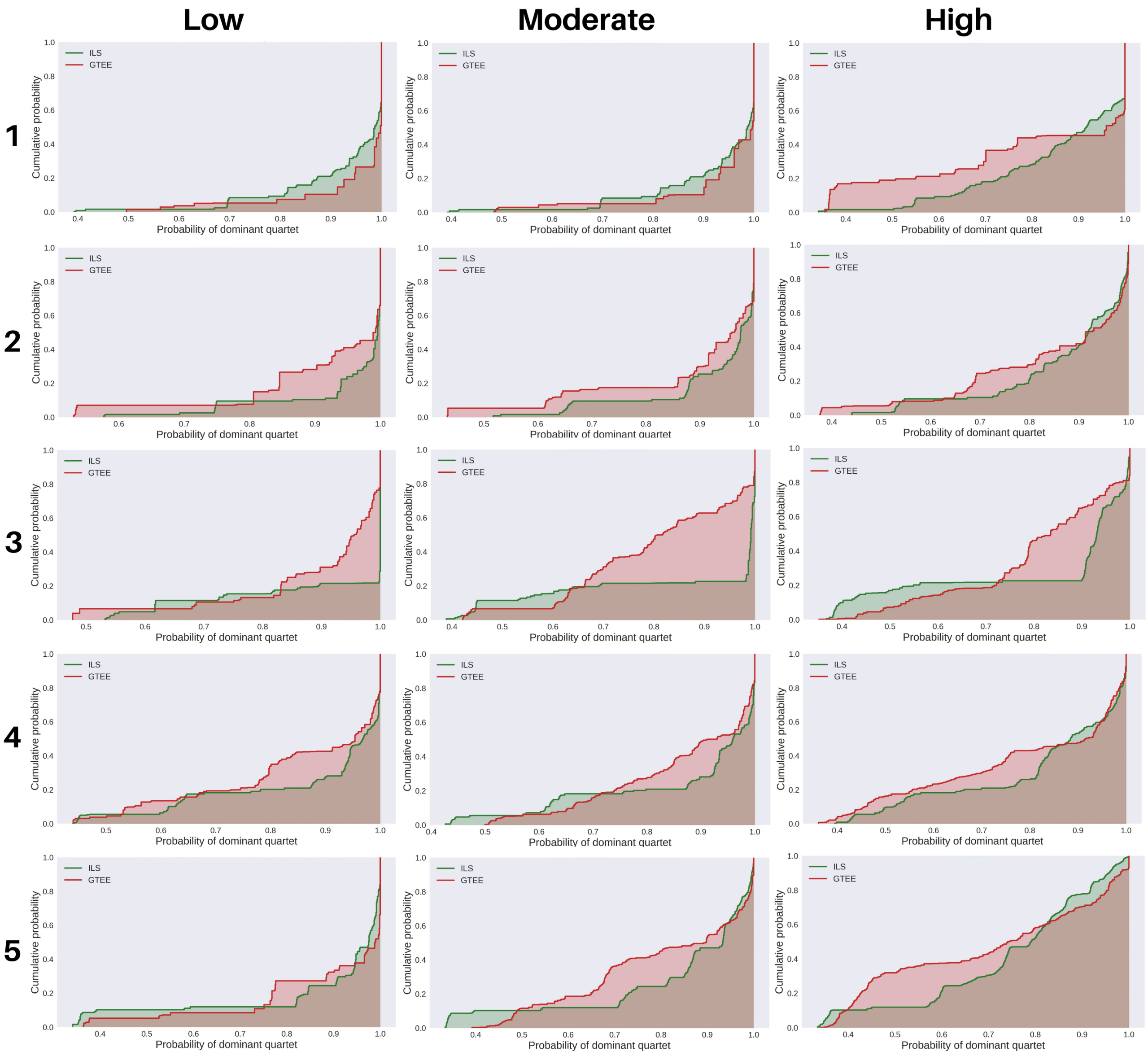


Figure 3: Empirical cumulative distribution functions of dominant quartet probabilities for the corresponding discordance levels (low, moderate and high) under matched ILS-only and GTEE-only conditions for 15-taxon dataset (replicate 1 to 5).

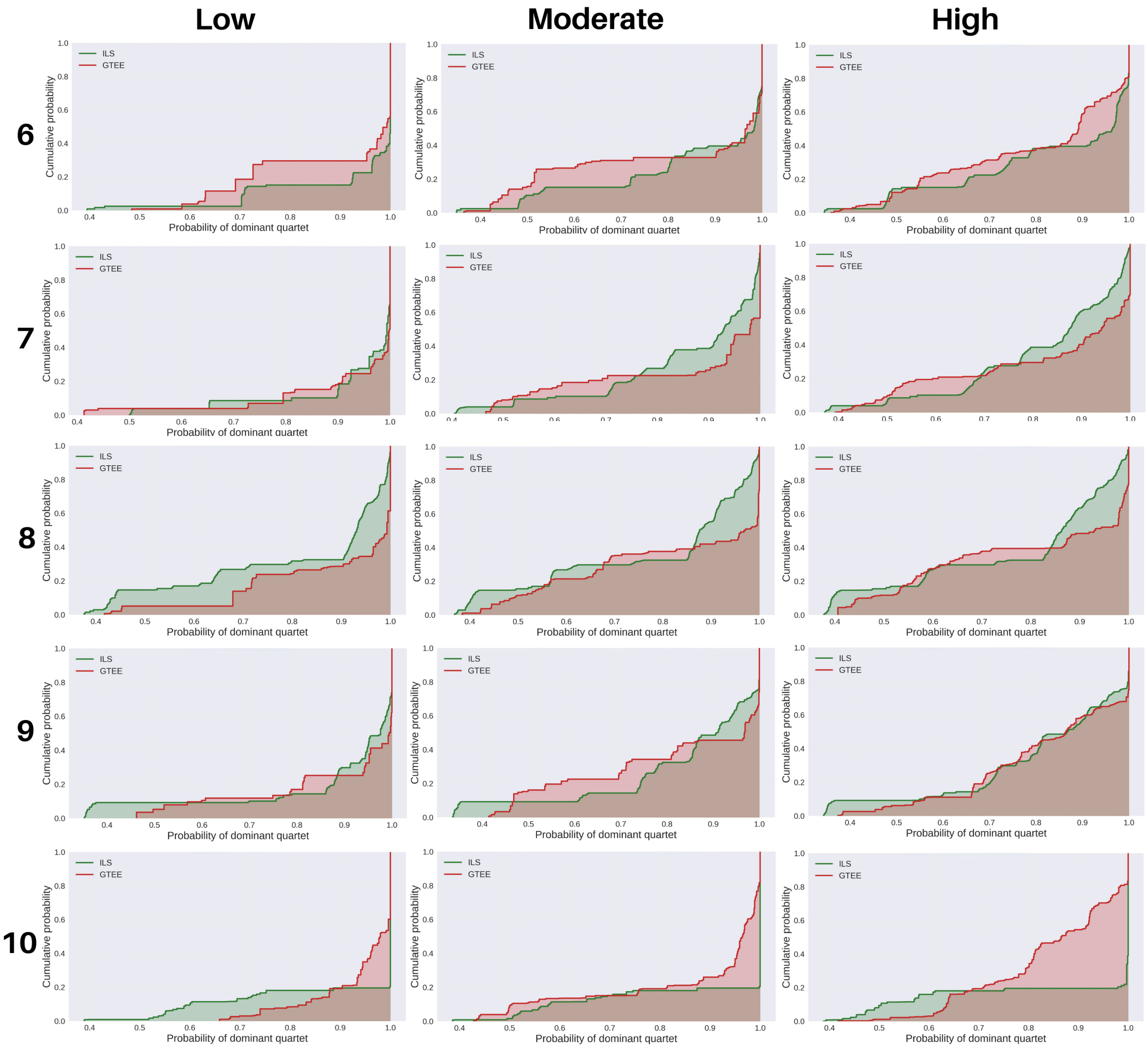


Figure 4: Empirical cumulative distribution functions of dominant quartet probabilities for the corresponding discordance levels (low, moderate and high) under matched ILS-only and GTEE-only conditions for 15-taxon dataset (replicate 6 to 10).

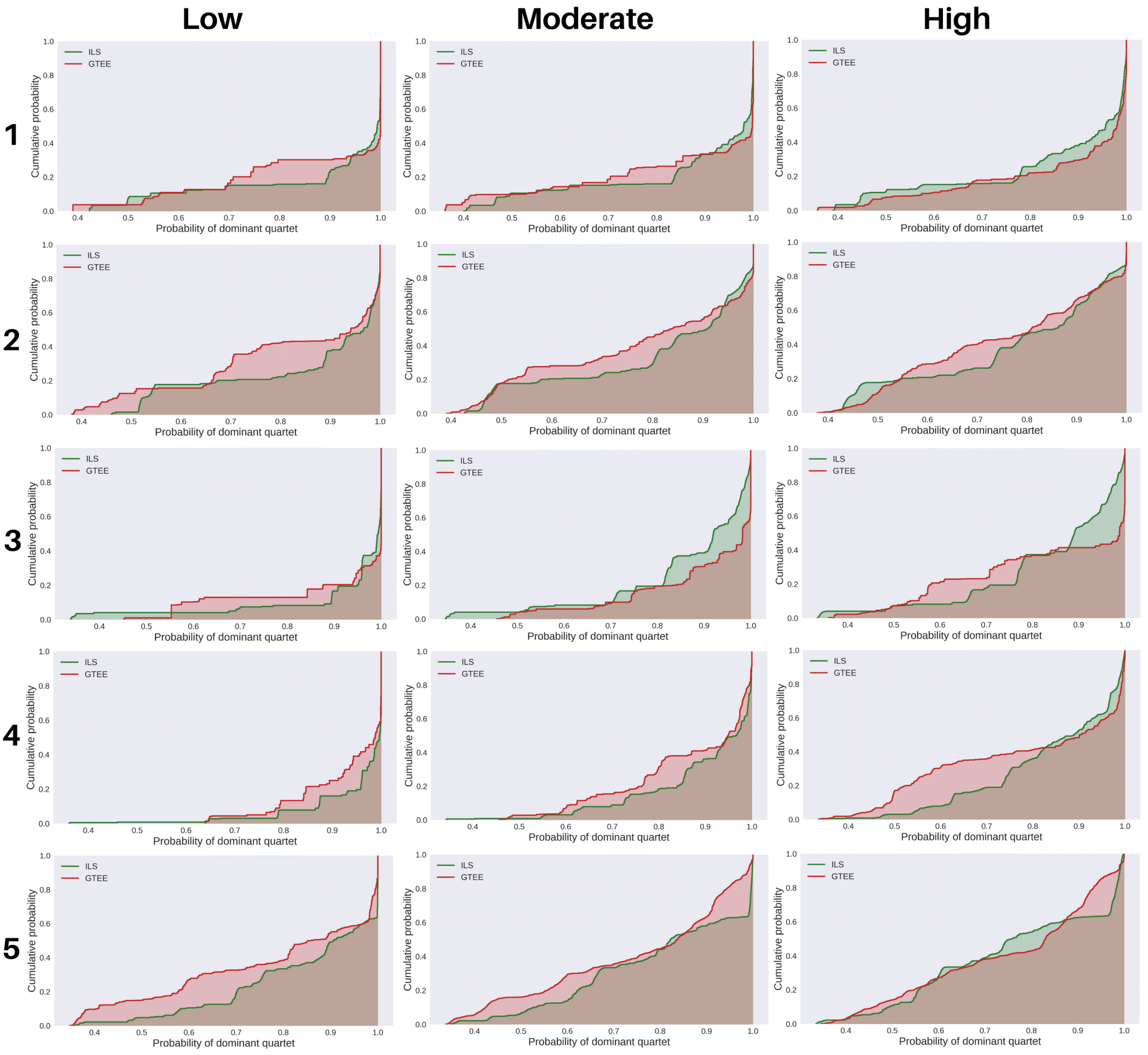


Figure 5: Empirical cumulative distribution functions of dominant quartet probabilities for the corresponding discordance levels (low, moderate and high) under matched ILS-only and GTEE-only conditions for 21-taxon dataset (replicates 1 to 5).

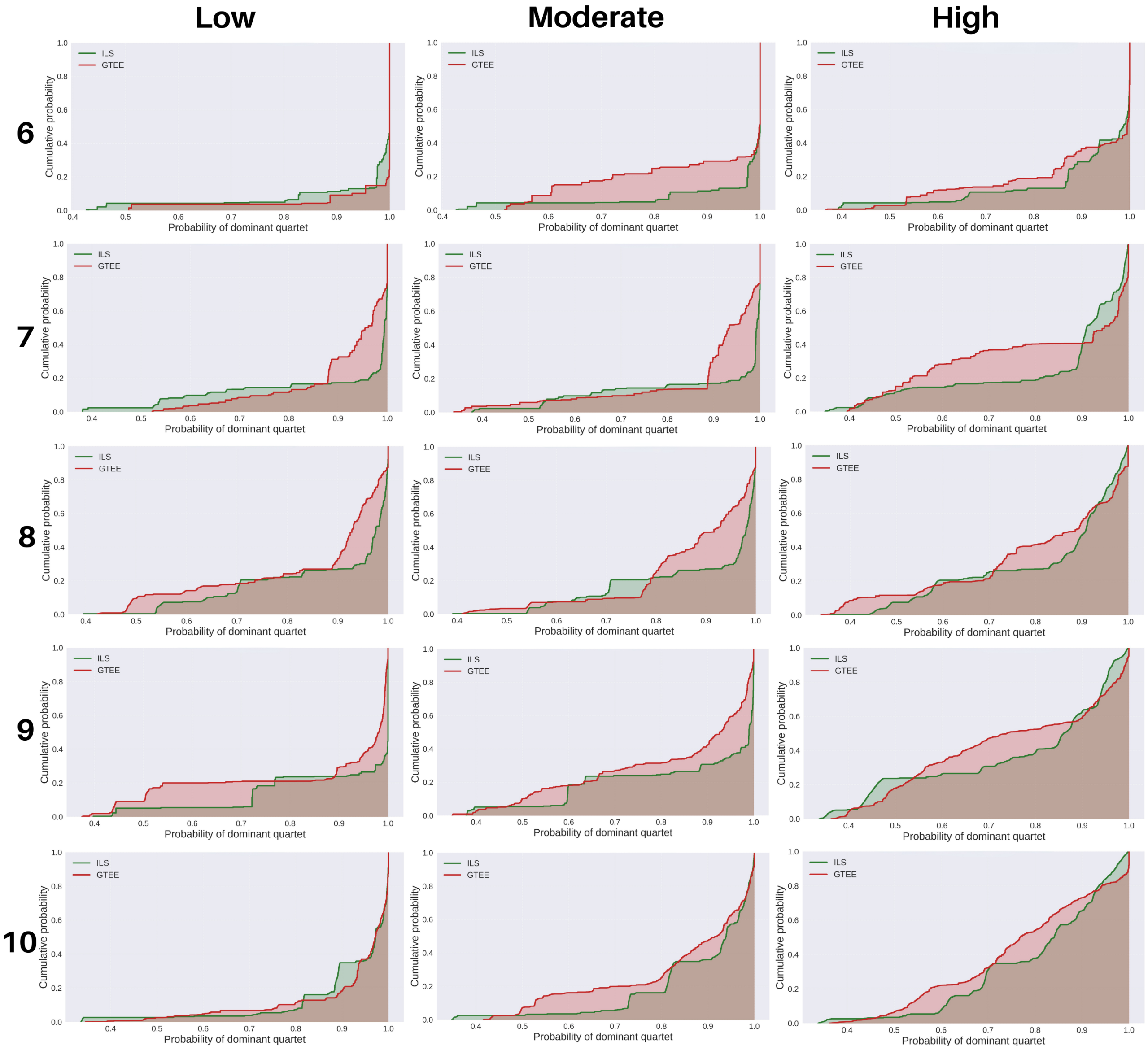


Figure 6: Empirical cumulative distribution functions of dominant quartet probabilities for the corresponding discordance levels (low, moderate and high) under matched ILS-only and GTEE-only conditions for 21-taxon dataset (replicates 6 to 10).

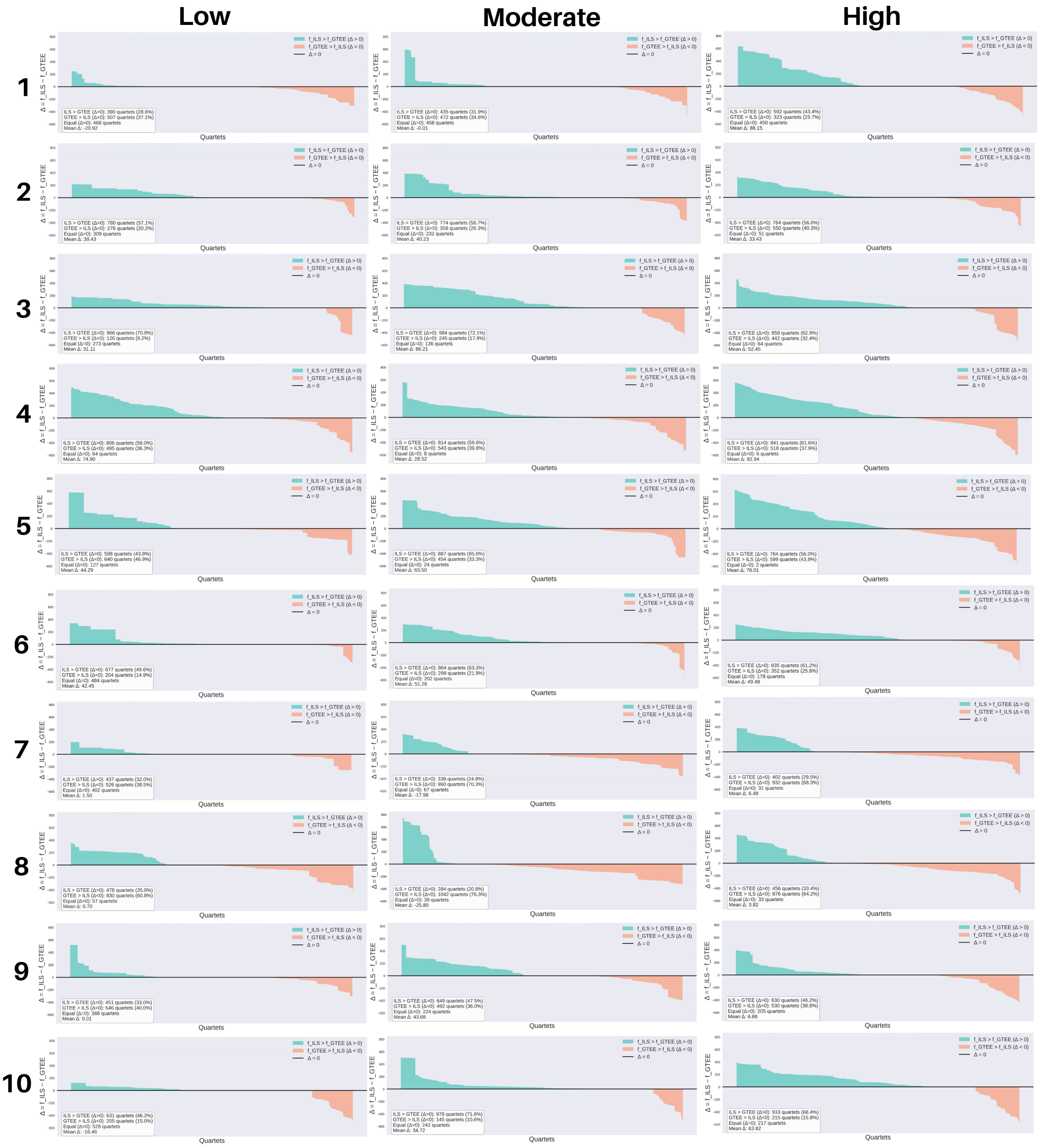

Figure 7: Sorted differences of true species-tree quartet ($q^*$) frequencies ($\Delta = \text{freq}_{\text{ILS}}(q^*) - \text{freq}_{\text{GTEE}}(q^*)$) across all quartets (total $\binom{15}{4} = 1365$ quartets) under low, moderate, and high discordance for 15-taxon dataset (all of 10 replicates)

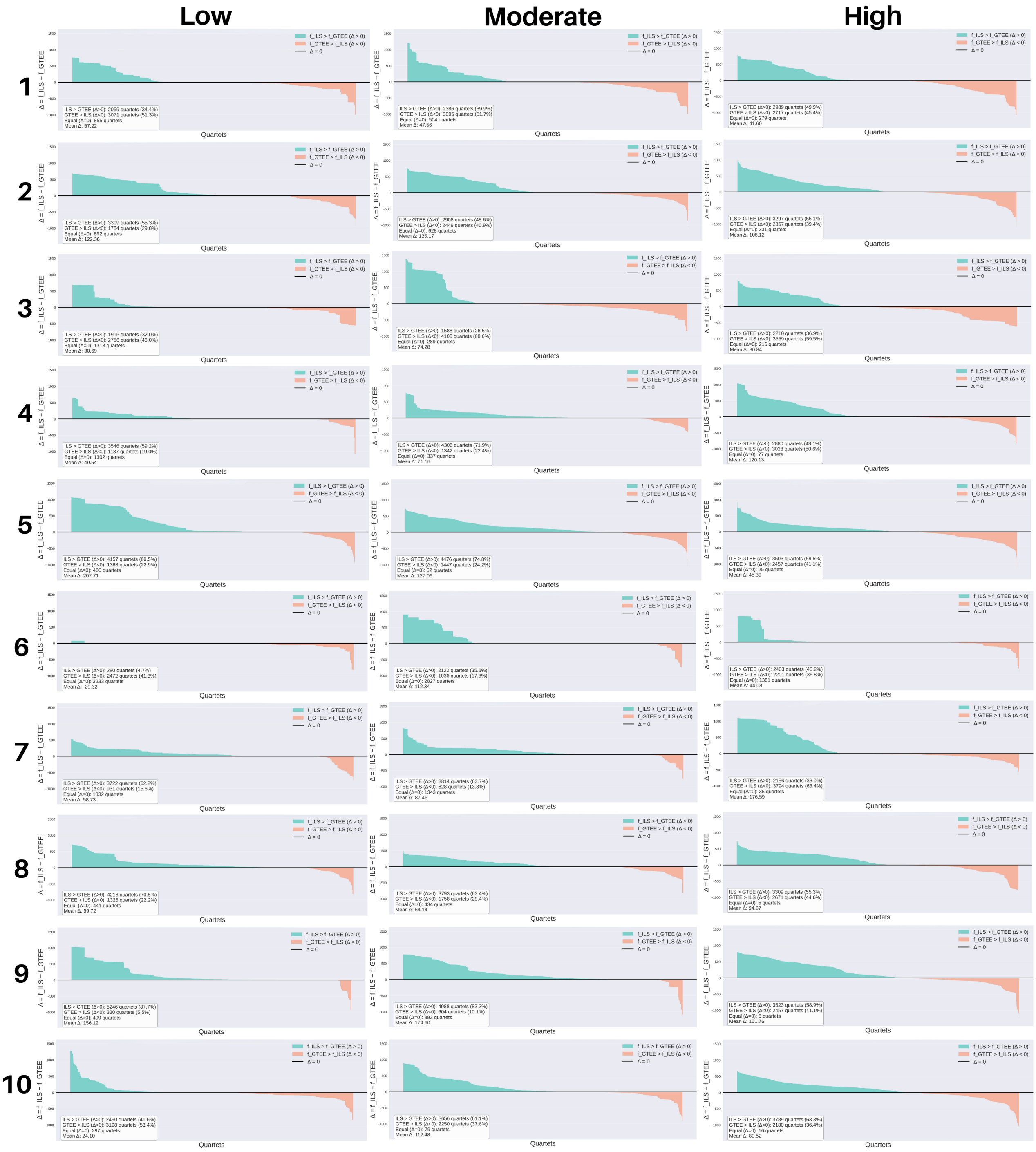


Figure 8: Sorted differences of true species-tree quartet ($q^*$) frequencies ($\Delta = \text{freq}_{\text{ILS}}(q^*) - \text{freq}_{\text{GTEE}}(q^*)$) across all quartets (total $\binom{21}{4} = 5985$ quartets) under low, moderate, and high discordance for 21-taxon dataset (all of 10 replicates)

### 3 Supplementary Tables

Table 4: Normalized RF distances for different methods under ILS-only and GTEE-only conditions across discordance levels and varying numbers of gene trees (25, 200, and 1000) for 15-taxon dataset. Values are reported as mean (standard error), computed across 10 replicates.

| Discordance | Method | 25 genes | | 200 genes | | 1000 genes | |
|---|---|---|---|---|---|---|---|
| | | ILS | GTEE | ILS | GTEE | ILS | GTEE |
| **Low** | ASTRAL | 0.083 (0.028) | 0.100 (0.027) | 0.017 (0.011) | 0.067 (0.024) | 0.008 (0.008) | 0.083 (0.028) |
| | WQFM | 0.083 (0.028) | 0.092 (0.029) | 0.017 (0.011) | 0.067 (0.024) | 0.008 (0.008) | 0.083 (0.028) |
| | PhyloNet | 0.083 (0.025) | 0.125 (0.028) | 0.058 (0.022) | 0.125 (0.028) | 0.050 (0.022) | 0.117 (0.025) |
| | Greedy | 0.083 (0.025) | 0.092 (0.029) | 0.025 (0.018) | 0.067 (0.024) | 0.008 (0.008) | 0.075 (0.029) |
| **Moderate** | ASTRAL | 0.075 (0.023) | 0.117 (0.040) | 0.025 (0.018) | 0.125 (0.028) | 0.008 (0.008) | 0.117 (0.036) |
| | WQFM | 0.083 (0.022) | 0.108 (0.035) | 0.025 (0.018) | 0.117 (0.031) | 0.008 (0.008) | 0.117 (0.036) |
| | PhyloNet | 0.075 (0.015) | 0.192 (0.039) | 0.058 (0.022) | 0.200 (0.028) | 0.067 (0.021) | 0.192 (0.031) |
| | Greedy | 0.067 (0.017) | 0.108 (0.035) | 0.025 (0.018) | 0.133 (0.033) | 0.042 (0.022) | 0.125 (0.036) |
| **High** | ASTRAL | 0.083 (0.022) | 0.192 (0.039) | 0.033 (0.018) | 0.208 (0.031) | 0.017 (0.017) | 0.208 (0.033) |
| | WQFM | 0.075 (0.019) | 0.217 (0.036) | 0.033 (0.018) | 0.208 (0.031) | 0.017 (0.017) | 0.200 (0.031) |
| | PhyloNet | 0.092 (0.015) | 0.283 (0.031) | 0.067 (0.021) | 0.283 (0.036) | 0.083 (0.025) | 0.275 (0.031) |
| | Greedy | 0.108 (0.018) | 0.225 (0.035) | 0.050 (0.022) | 0.217 (0.033) | 0.058 (0.028) | 0.217 (0.033) |

Table 5: Normalized RF distances for different methods under ILS-only and GTEE-only conditions across discordance levels and varying numbers of gene trees (100, 400, and 2000) for 21-taxon dataset. Values are reported as mean (standard error), computed across 10 replicates.

| Discordance | Method | 100 genes | | 400 genes | | 2000 genes | |
|---|---|---|---|---|---|---|---|
| | | ILS | GTEE | ILS | GTEE | ILS | GTEE |
| **Low** | ASTRAL | 0.011 (0.007) | 0.111 (0.031) | 0.006 (0.006) | 0.100 (0.023) | 0.006 (0.006) | 0.106 (0.024) |
| | WQFM | 0.011 (0.007) | 0.111 (0.027) | 0.006 (0.006) | 0.106 (0.024) | 0.006 (0.006) | 0.100 (0.022) |
| | PhyloNet | 0.017 (0.008) | 0.111 (0.020) | 0.017 (0.008) | 0.100 (0.020) | 0.017 (0.008) | 0.100 (0.020) |
| | Greedy | 0.011 (0.007) | 0.094 (0.020) | 0.006 (0.006) | 0.100 (0.023) | 0.011 (0.007) | 0.111 (0.027) |
| **Moderate** | ASTRAL | 0.033 (0.012) | 0.122 (0.032) | 0.017 (0.012) | 0.117 (0.031) | 0.011 (0.007) | 0.122 (0.034) |
| | WQFM | 0.028 (0.009) | 0.128 (0.035) | 0.017 (0.012) | 0.128 (0.032) | 0.011 (0.007) | 0.133 (0.034) |
| | PhyloNet | 0.028 (0.009) | 0.150 (0.026) | 0.039 (0.014) | 0.150 (0.029) | 0.028 (0.009) | 0.144 (0.025) |
| | Greedy | 0.028 (0.009) | 0.122 (0.032) | 0.017 (0.012) | 0.128 (0.032) | 0.011 (0.007) | 0.128 (0.034) |
| **High** | ASTRAL | 0.039 (0.012) | 0.189 (0.024) | 0.033 (0.012) | 0.178 (0.022) | 0.011 (0.007) | 0.172 (0.023) |
| | WQFM | 0.033 (0.012) | 0.189 (0.022) | 0.033 (0.012) | 0.183 (0.022) | 0.011 (0.007) | 0.178 (0.022) |
| | PhyloNet | 0.061 (0.013) | 0.261 (0.023) | 0.056 (0.014) | 0.256 (0.028) | 0.061 (0.015) | 0.267 (0.026) |
| | Greedy | 0.033 (0.009) | 0.194 (0.021) | 0.044 (0.014) | 0.178 (0.019) | 0.017 (0.008) | 0.183 (0.022) |

Table 6: Summary statistics of quartet distributions and species-tree neighborhood analyses under ILS-only and GTEE-only conditions across discordance levels and replicates for the **15-taxon** dataset. For each replicate and discordance level (low, moderate, high), the table reports the percentage of true species-tree quartets that are dominant (out of 1365 total quartets) in the gene-tree–induced quartet distribution, the skewness and kurtosis, the average entropy and dispersion of dominant quartet probability distributions, and the number of neighboring candidate species-tree topologies with quartet scores exceeding that of the true species tree.

| **Setup** | | **Condition** | **%Dominant** | **Skewness** | **Entropy** | **Dispersion** | **#neighbors score >** |
|---|---|---|---|---|---|---|---|
| **Replicate** | **Discordance** | | **Quartets (1365)** | | | | **true ST score** |
| R1 | low | ILS | 100.0000 | -2.4897 | 0.2120 | 0.0659 | 0 |
| | | GTEE | 100.0000 | -3.1296 | 0.1384 | 0.0450 | 0 |
| | moderate | ILS | 100.0000 | -2.4897 | 0.2120 | 0.0659 | 0 |
| | | GTEE | 94.8718 | -3.0545 | 0.1662 | 0.0538 | 9 |
| | high | ILS | 100.0000 | -1.1279 | 0.3902 | 0.1390 | 0 |
| | | GTEE | 86.6667 | -0.7714 | 0.4266 | 0.2025 | 31 |
| R2 | low | ILS | 100.0000 | -2.7269 | 0.1417 | 0.0412 | 0 |
| | | GTEE | 100.0000 | -2.0071 | 0.2311 | 0.0807 | 0 |
| | moderate | ILS | 100.0000 | -2.1767 | 0.2213 | 0.0672 | 0 |
| | | GTEE | 100.0000 | -1.7371 | 0.2967 | 0.1074 | 0 |
| | high | ILS | 100.0000 | -1.4805 | 0.3730 | 0.1235 | 0 |
| | | GTEE | 94.6520 | -1.2137 | 0.3874 | 0.1477 | 4 |
| R3 | low | ILS | 100.0000 | -1.8713 | 0.1725 | 0.0670 | 0 |
| | | GTEE | 97.3626 | -1.9794 | 0.2778 | 0.0974 | 0 |
| | moderate | ILS | 100.0000 | -1.5273 | 0.2500 | 0.1100 | 0 |
| | | GTEE | 96.0440 | -0.5189 | 0.4984 | 0.1946 | 0 |
| | high | ILS | 100.0000 | -1.3370 | 0.4003 | 0.1621 | 0 |
| | | GTEE | 92.4542 | -0.7887 | 0.5065 | 0.1894 | 115 |
| R4 | low | ILS | 100.0000 | -1.6108 | 0.2948 | 0.1064 | 0 |
| | | GTEE | 92.2344 | -1.0427 | 0.3734 | 0.1424 | 50 |
| | moderate | ILS | 100.0000 | -1.5752 | 0.3108 | 0.1120 | 0 |
| | | GTEE | 95.4579 | -0.9861 | 0.3683 | 0.1303 | 4 |
| | high | ILS | 100.0000 | -1.0649 | 0.4480 | 0.1668 | 0 |
| | | GTEE | 76.7766 | -0.6854 | 0.4793 | 0.2025 | 56 |
| R5 | low | ILS | 99.1941 | -2.1152 | 0.2885 | 0.1124 | 7 |
| | | GTEE | 86.0806 | -1.8595 | 0.2772 | 0.1064 | 65 |
| | moderate | ILS | 94.0659 | -1.6411 | 0.4221 | 0.1587 | 19 |
| | | GTEE | 86.3736 | -0.5263 | 0.4905 | 0.1950 | 32 |
| | high | ILS | 91.4286 | -0.8333 | 0.6521 | 0.2540 | 42 |
| | | GTEE | 76.2637 | -0.1800 | 0.6552 | 0.2975 | 137 |
| R6 | low | ILS | 100.0000 | -2.4595 | 0.1758 | 0.0616 | 0 |
| | | GTEE | 99.1209 | -1.0464 | 0.2716 | 0.1034 | 1 |
| | moderate | ILS | 100.0000 | -1.2814 | 0.3515 | 0.1369 | 0 |
| | | GTEE | 90.4762 | -0.8224 | 0.3867 | 0.1757 | 8 |
| | high | ILS | 100.0000 | -1.0062 | 0.4093 | 0.1612 | 0 |
| | | GTEE | 93.8462 | -0.7020 | 0.4681 | 0.1982 | 8 |
| R7 | low | ILS | 100.0000 | -2.6443 | 0.1837 | 0.0595 | 0 |
| | | GTEE | 99.1209 | -2.8836 | 0.1598 | 0.0607 | 2 |
| | moderate | ILS | 100.0000 | -1.3694 | 0.3928 | 0.1366 | 0 |
| | | GTEE | 97.8022 | -1.3579 | 0.2706 | 0.1185 | 0 |
| | high | ILS | 100.0000 | -1.0663 | 0.4860 | 0.1734 | 0 |
| | | GTEE | 85.0549 | -1.0183 | 0.3746 | 0.1570 | 17 |
| R8 | low | ILS | 100.0000 | -1.0680 | 0.4353 | 0.1725 | 0 |
| | | GTEE | 84.9084 | -1.5215 | 0.2666 | 0.1022 | 73 |
| | moderate | ILS | 100.0000 | -0.8762 | 0.5216 | 0.2095 | 0 |
| | | GTEE | 91.7949 | -0.7288 | 0.3920 | 0.1714 | 11 |
| | high | ILS | 100.0000 | -0.8585 | 0.5488 | 0.2188 | 0 |
| | | GTEE | 88.8645 | -0.5730 | 0.4481 | 0.1950 | 15 |
| R9 | low | ILS | 100.0000 | -2.3008 | 0.2758 | 0.1045 | 0 |
| | | GTEE | 93.4066 | -1.7720 | 0.2415 | 0.0932 | 16 |
| | moderate | ILS | 96.7766 | -1.5196 | 0.4259 | 0.1617 | 0 |
| | | GTEE | 89.1575 | -0.7763 | 0.4051 | 0.1747 | 53 |
| | high | ILS | 99.3407 | -1.1576 | 0.4859 | 0.1865 | 0 |
| | | GTEE | 93.1136 | -0.7640 | 0.4570 | 0.1742 | 46 |
| R10 | low | ILS | 100.0000 | -1.8947 | 0.1718 | 0.0714 | 0 |
| | | GTEE | 100.0000 | -1.8589 | 0.1853 | 0.0549 | 0 |
| | moderate | ILS | 100.0000 | -1.8491 | 0.1776 | 0.0761 | 0 |
| | | GTEE | 100.0000 | -1.6900 | 0.2898 | 0.1108 | 0 |
| | high | ILS | 100.0000 | -1.6681 | 0.2008 | 0.0926 | 0 |
| | | GTEE | 100.0000 | -0.6179 | 0.4490 | 0.1564 | 0 |

Table 7: Summary statistics of quartet distributions and species-tree neighborhood analyses under ILS-only and GTEE-only conditions across discordance levels and replicates for the **21-taxon** dataset. For each replicate and discordance level (low, moderate, high), the table reports the percentage of true species-tree quartets that are dominant (out of 5985 total quartets) in the gene-tree–induced quartet distribution, the skewness and kurtosis, the average entropy and dispersion of dominant quartet probability distributions, and the number of neighboring candidate species-tree topologies with quartet scores exceeding that of the true species tree.

| Setup | | Condition | %Dominant | Skewness | Entropy | Dispersion | #neighbors score > |
|---|---|---|---|---|---|---|---|
| Replicate | Discordance | | Quartets (5985) | | | | true ST score |
| R1 | low | ILS | 100.0000 | -1.9407 | 0.2362 | 0.0902 | 0 |
| | | GTEE | 92.7820 | -1.3765 | 0.2585 | 0.1121 | 4 |
| | moderate | ILS | 100.0000 | -1.6721 | 0.2993 | 0.1137 | 0 |
| | | GTEE | 94.1186 | -1.4244 | 0.2988 | 0.1297 | 10 |
| | high | ILS | 100.0000 | -1.4262 | 0.3598 | 0.1372 | 0 |
| | | GTEE | 89.8413 | -1.6185 | 0.2931 | 0.1132 | 72 |
| R2 | low | ILS | 100.0000 | -1.2612 | 0.3489 | 0.1298 | 0 |
| | | GTEE | 91.7126 | -0.7853 | 0.4169 | 0.1792 | 23 |
| | moderate | ILS | 100.0000 | -0.9576 | 0.4716 | 0.1795 | 0 |
| | | GTEE | 89.3901 | -0.4850 | 0.5077 | 0.2156 | 108 |
| | high | ILS | 100.0000 | -0.7199 | 0.5466 | 0.2137 | 0 |
| | | GTEE | 83.2581 | -0.2775 | 0.5707 | 0.2344 | 113 |
| R3 | low | ILS | 97.6107 | -3.4234 | 0.1726 | 0.0597 | 3 |
| | | GTEE | 99.4319 | -1.9824 | 0.1881 | 0.0743 | 8 |
| | moderate | ILS | 96.5246 | -1.7442 | 0.3858 | 0.1324 | 3 |
| | | GTEE | 81.1529 | -1.7369 | 0.2513 | 0.0920 | 888 |
| | high | ILS | 97.9616 | -1.3829 | 0.4495 | 0.1592 | 3 |
| | | GTEE | 90.8104 | -0.8380 | 0.3566 | 0.1571 | 49 |
| R4 | low | ILS | 100.0000 | -3.4892 | 0.1499 | 0.0436 | 0 |
| | | GTEE | 99.4319 | -1.6131 | 0.2125 | 0.0655 | 1 |
| | moderate | ILS | 99.4319 | -1.6502 | 0.3063 | 0.0985 | 3 |
| | | GTEE | 99.4319 | -1.0183 | 0.3696 | 0.1305 | 10 |
| | high | ILS | 99.4319 | -0.8676 | 0.4611 | 0.1604 | 3 |
| | | GTEE | 94.5196 | -0.5155 | 0.4895 | 0.2085 | 51 |
| R5 | low | ILS | 100.0000 | -1.0606 | 0.3963 | 0.1447 | 0 |
| | | GTEE | 82.4060 | -0.6955 | 0.4829 | 0.2106 | 247 |
| | moderate | ILS | 100.0000 | -0.5432 | 0.4987 | 0.1948 | 0 |
| | | GTEE | 87.7694 | -0.6043 | 0.5828 | 0.2419 | 115 |
| | high | ILS | 99.5656 | -0.2751 | 0.5787 | 0.2350 | 0 |
| | | GTEE | 91.3450 | -0.5214 | 0.5920 | 0.2410 | 430 |
| R6 | low | ILS | 100.0000 | -3.5767 | 0.1204 | 0.0423 | 0 |
| | | GTEE | 100.0000 | -4.3010 | 0.0721 | 0.0276 | 0 |
| | moderate | ILS | 100.0000 | -3.5767 | 0.1204 | 0.0423 | 0 |
| | | GTEE | 100.0000 | -1.3080 | 0.2310 | 0.0985 | 0 |
| | high | ILS | 100.0000 | -2.3375 | 0.2473 | 0.0850 | 0 |
| | | GTEE | 99.4319 | -1.5359 | 0.2627 | 0.1041 | 5 |
| R7 | low | ILS | 100.0000 | -2.1341 | 0.1895 | 0.0745 | 0 |
| | | GTEE | 93.8513 | -1.8727 | 0.2562 | 0.0831 | 50 |
| | moderate | ILS | 100.0000 | -2.1020 | 0.1940 | 0.0763 | 0 |
| | | GTEE | 91.8463 | -2.3289 | 0.2956 | 0.1038 | 63 |
| | high | ILS | 100.0000 | -1.6016 | 0.3960 | 0.1444 | 0 |
| | | GTEE | 83.6424 | -0.6106 | 0.4383 | 0.1937 | 81 |
| R8 | low | ILS | 100.0000 | -1.4351 | 0.2874 | 0.0995 | 0 |
| | | GTEE | 96.0902 | -1.3761 | 0.3874 | 0.1394 | 17 |
| | moderate | ILS | 100.0000 | -1.4509 | 0.2784 | 0.0958 | 0 |
| | | GTEE | 100.0000 | -1.5067 | 0.3772 | 0.1279 | 0 |
| | high | ILS | 100.0000 | -1.0688 | 0.4683 | 0.1678 | 0 |
| | | GTEE | 93.0326 | -0.9203 | 0.4960 | 0.1977 | 51 |
| R9 | low | ILS | 100.0000 | -1.8825 | 0.2151 | 0.0821 | 0 |
| | | GTEE | 87.0008 | -1.4040 | 0.3102 | 0.1301 | 61 |
| | moderate | ILS | 100.0000 | -1.3386 | 0.3048 | 0.1227 | 0 |
| | | GTEE | 83.9933 | -0.9933 | 0.4246 | 0.1687 | 134 |
| | high | ILS | 99.9833 | -0.7859 | 0.5754 | 0.2320 | 0 |
| | | GTEE | 79.1145 | -0.1881 | 0.5878 | 0.2565 | 252 |
| R10 | low | ILS | 98.9641 | -3.0493 | 0.2478 | 0.0773 | 0 |
| | | GTEE | 95.0209 | -2.5980 | 0.2284 | 0.0730 | 15 |
| | moderate | ILS | 100.0000 | -1.9493 | 0.3535 | 0.1134 | 0 |
| | | GTEE | 95.5221 | -1.1296 | 0.4040 | 0.1541 | 16 |
| | high | ILS | 99.8997 | -0.8808 | 0.5599 | 0.1972 | 0 |
| | | GTEE | 96.1069 | -0.2886 | 0.5857 | 0.2319 | 6 |

### References

[1] Diego Mallo, Leonardo de Oliveira Martins, and David Posada. “SimPhy: phylogenomic simulation of gene, locus, and species trees”. In: *Systematic Biology* 65.2 (2016), pp. 334–344.

[2] Nhan Ly-Trong et al. “AliSim: a fast and versatile phylogenetic sequence simulator for the genomic era”. In: *Molecular Biology and Evolution* 39.5 (2022), msac092.

[3] Thomas KF Wong et al. “IQ-TREE 3: phylogenomic inference software using complex evolutionary models”. In: *Molecular Biology and Evolution* 43.5 (2026), msag117.

[4] Chao Zhang et al. “ASTRAL-III: polynomial time species tree reconstruction from partially resolved gene trees”. In: *BMC Bioinformatics* 19.6 (2018), p. 153.
